# Essential function of *femaleless* in female gametogenesis controls gene drive spread in *Anopheles gambiae*

**DOI:** 10.64898/2026.08.22.746406

**Authors:** B. Fasulo, W. Garrood, J. Philpott, L. Marston, K. Willis, N. Kranjc, A. Strampelli, A. Burt, F. Bernardini, A. Crisanti

## Abstract

Insecticide resistance in mosquito vectors and antimalarial drug resistance in parasites threaten progress towards malaria elimination, prompting the development of alternative control strategies such as CRISPR-Cas9 gene drives. The sex determination gene *femaleless* (*fle*, AGAP013051), which is required for female development in *Anopheles gambiae,* is a promising target for population-suppression approaches aimed at disrupting female-specific genes that affect fertility or viability. However, its functions beyond sex determination remain unknown.

Here, we engineered homing gene drives targeting *fle* and employed germline promoters with distinct temporal expression profiles, early-acting *zero population growth* (*zpg,* AGAP006241) and late-acting *sporulation defective 11* (*spo11*, AGAP010898), to modulate Cas9 activity. The *zpg*-driven system achieved up to 98% transmission through males but caused complete sterility in hemizygous females due to early biallelic disruption of *fle* during germline development. Delaying *cas9* expression with the *spo11* promoter partially restored female fertility, although female transmission remained close to Mendelian levels (59%). These results reveal an essential role for *fle* in female gametogenesis in addition to its established function in sex determination. Population modelling predicts that releasing *zpg*-drive males at 16.9% of the wild-type male population could reduce female abundance by 95% within 36 generations. Collectively, our findings reveal a previously unrecognised reproductive function of *fle* that limits gene-drive spread and provide important insights for the design of vector-control strategies targeting genes with essential germline functions.

## Introduction

Malaria remains a major global health issue, resulting in 610,000 deaths and 282 million estimated cases in 2024 (WHO, 2025). Mosquitoes of the *Anopheles gambiae* complex are the primary vector of *Plasmodium* parasites in sub-Saharan Africa, where the disease burden is highest. Currently, efforts to control and eradicate malaria rely heavily on insecticides and antimalarial drugs; however, the widespread emergence of resistance is hindering their effectiveness (Ranson and Lissenden, 2016; Hancock *et al*., 2020; Wilson *et al*., 2020), creating an urgent need for complementary control technology.

Genetic control strategies are emerging as promising tools for malaria vector management because they can spread engineered traits through mosquito populations while reducing their reproductive capacity. Such approaches can be broadly divided into non-driving systems, which are inherited according to Mendelian principles and transmitted to approximately 50% of the offspring of an heterozygous parent, and gene drive systems, which bias inheritance to achieve super-Mendelian transmission rates greater than 50%, thereby facilitating their rapid spread through target populations (Alphey, 2013).

Advances in CRISPR–Cas9 genome engineering have accelerated the development of highly efficient gene drive systems in *An. gambiae* (Gantz *et al*., 2015; Hammond *et al*., 2016; Kyrou *et al*., 2018; Adolfi *et al*., 2020; Carballar-Lejarazú *et al*., 2020; Simoni *et al*., 2020; Xu *et al*., 2025). Most of these systems rely on a mechanism known as homing, in which the endonuclease induces a double-strand break at the target locus in germline cells. Repair of the cut through Homology-Directed Repair (HDR), using the drive-containing chromosome as a template, results in copying the gene drive onto the homologous chromosome. This process converts heterozygotes into homozygotes and promotes super-Mendelian inheritance in future generations.

The efficient spread of homing gene drives through populations has enabled the development of population-suppression strategies targeting genes essential for female development and fertility. In *An. gambiae*, targeting the sex-determination gene *doublesex* (*dsx*) provided proof of principle that disrupting female reproductive function can impose a substantial genetic load on mosquito populations, leading to marked reductions in female fertility and population collapse under laboratory conditions (Hammond *et al*., 2016; Kyrou *et al*., 2018; Hammond *et al*., 2021b; Strampelli *et al*., 2025). Sex-specific alternative splicing of *dsx* generates two isoforms: *dsxF*, which is required for female sexual development and fertility, and *dsxM*, which directs male sexual development and function. Disruption of the female-specific *dsxF* isoform results in sterile females and males largely unaffected, providing a highly sex-specific mechanism for population suppression.

More recently, genetic strategies targeting *dsx* in other mosquito species have further validated this gene as an attractive target for genetic population-suppression approaches (Xu *et al*., 2025). Despite these findings, relatively few genes essential for female viability or fertility have been systematically explored as candidate targets for vector control, and even fewer have been functionally characterised in sufficient mechanistic detail to predict how their disruption may influence both reproductive fitness and gene-drive dynamics.

*femaleless* (*fle*; AGAP013051) is a key regulator of female development in *An. gambiae* (Krzywinska *et al*., 2021). Acting upstream of *dsx* and *fruitless* (*fru*), *fle* directs female-specific splicing of both genes and suppresses Dosage Compensation (DC) in females (Kalita *et al*., 2023; Krzywinska *et al*., 2023). Although expressed in both sexes, Fle is primarily required in females, where its loss disrupts normal development.

Functional studies have demonstrated the critical role of Fle during female development. Transient embryonic knockdown causes female-specific lethality, whereas stable microRNA-mediated silencing produces a spectrum of phenotypes ranging from partial masculinization to complete lethality (Krzywinska *et al*., 2021). While these findings initially suggested a dose-dependent requirement for Fle, subsequent work revealed that the gene is haplosufficient. Using a split gene-drive system, Smidler *et al*. showed that females retaining a single functional *fle* allele following maternal Cas9-induced mutagenesis remained viable and fertile, indicating that one intact copy is sufficient to support normal female development (Smidler *et al*., 2023).

Together, these characteristics - female-specific, essentiality, haplosufficiency, and a central regulatory role within the sex-determination pathway - identify *fle* as a promising target for genetic population-suppression strategies.

Current understanding of Fle function derives largely from studies of embryonic development and sex determination, leaving potential roles in adult female reproduction unexplored. Determining whether *fle* contributes to female reproductive processes is important not only for assessing its suitability as a target for genetic control but also for understanding how previously unrecognised gene functions may influence the performance and spread of suppression gene drives.

To investigate the potential of *fle* as a gene-drive target, we developed a multiplex homing gene drive using the early-acting germline promoter *zero population growth* (*zpg*; AGAP006241) to drive *cas9* expression (Thailayil *et al*., 2011; Kyrou *et al*., 2018; Hammond *et al*., 2021a). Although this system achieved near-complete drive transmission in males, it unexpectedly induced complete sterility in heterozygous females despite previous evidence that *fle* is haplosufficient for female development (Smidler *et al*., 2023).

This observation suggested that *fle* may perform an additional essential function during female reproduction that is sensitive to the timing of gene disruption, prompting us to investigate the mechanism underlying the dominant sterility phenotype. We hypothesised that early *zpg-*driven *cas9* expression disrupts *fle* activity during a critical window of female germline development, before oocyte differentiation. To test whether delaying *cas9* expression could preserve female fertility while maintaining drive efficiency, we engineered a second drive system using the later-acting meiotic promoter *sporulation defective 11* (*spo11/*AGAP010898) (Terradas *et al*., 2021). By comparing the performance of these two systems, we sought to define the temporal requirements for Fle function during female gametogenesis and evaluate whether this locus can support efficient and sustainable gene-drive transmission.

## Results

### Generation and characterisation of a *femaleless* knockout strain in *Anopheles gambiae*

To evaluate *femaleless* as a gene drive target, we first generated a knockout allele with an integrated docking site for subsequent insertion of a gene drive cassette (Krzywinska *et al*., 2021). Using two guide RNAs (gRNAs), *13051T14* (*T14*) and *13051T1* (*T1*), targeting highly conserved regions (Kranjc *et al*., 2021) within *fle* second RNA recognition motif (RRM; Fig S1A), we replaced a 74 bp fragment with a GFP marker cassette (*3xP3::GFP*) flanked by two *attP* recombination sites (Fig. 1A).

**Figure 1.**
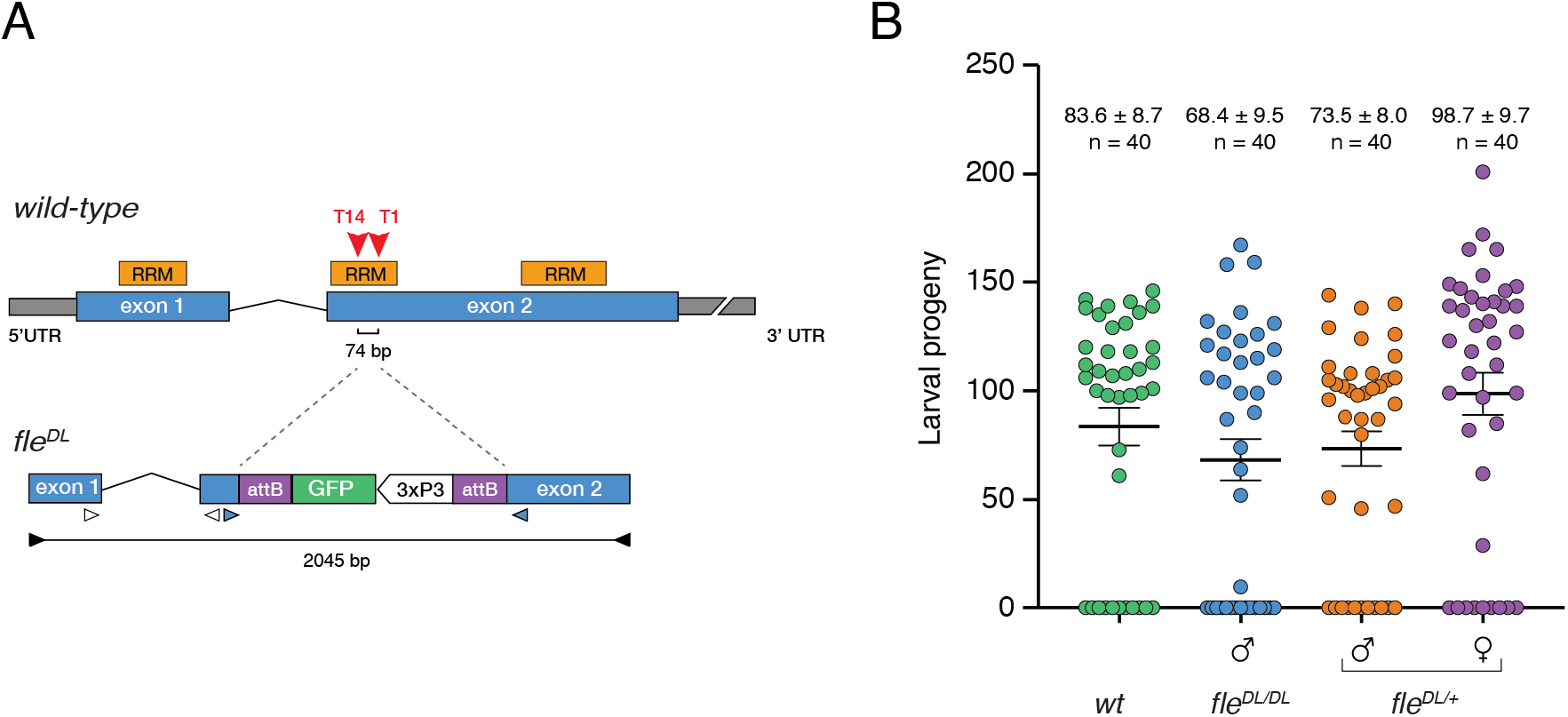
Generations and characterisation of the *femaleless* docking line *fle^DL^*. (**A**) Top: Schematic of the *femaleless* (*fle*) gene structure, showing two exons (blue boxes) and untranslated regions (grey). The gene contains three RNA recognition motifs (RRMs), one in exon 1 and two in exon 2. Guide RNA target sites *T1* and *T14* (arrowheads) flank a 74 bp region within the second RRM. Bottom: Strategy for generating the *fle^DL^*allele. The donor construct contains a *3XP3::GFP* marker cassette flanked by *attP* recombination sites, with left (LHA) and right (RHA) homology arms for homologous integration. Following integration, the 74 bp deletion is replaced by the *GFP* cassette, yielding a 2045 bp diagnostic PCR amplicon (primers *WG162/WG163*), compared with 722 bp in the wild-type. The white triangles represent primers used for the qPCR (*WG134/WG135*), and the blue triangles represent primers used for the RT-PCR (*WG128/WG129*). (**B**) Fertility assessment of *fle^DL^*males and females crossed to wild-type. Larval progeny counts from individual crosses show that heterozygous females (*fle^DL/+^*) have fertility comparable to wild-type (*wt*), confirming haplosufficiency. Homozygous and heterozygous males show normal or enhanced fertility. Each point represents one genetic cross; bars indicate mean ± s.e.m. No significant differences were detected (Kruskal-Wallis test with Dunn’s multiple comparisons versus wild-type; p = 0.3689 for *fle^DL/+^*females, p = 0.8558 for *fle^DL/DL^* males, and p = 0.6550 for *fle^DL/+^* males. n = 40 for all groups.

The resulting *fle^Docking^ ^Line^* (*fle^DL^*) allele encodes a truncated, out-of-frame protein of 167 amino acids compared to the wild-type 420 amino acids (Krzywinska *et al*., 2021). The cassette insertion in the deleted region was confirmed with diagnostic PCR using primers spanning the insertion site (Fig. S1B). Sequencing of the amplicon also confirmed the deletion of the 74 bp and its replacement with the GFP cassette (Data Source).

To assess the *fle^DL^* knockout phenotype, we intercrossed hemizygous mosquitoes. At the pupal stage, *fle^DL/+^* progeny showed the expected 1:1 male-to-female ratio, whereas all *fle^DL/DL^*pupae were morphologically male. PCR amplification of the Y-linked *yob* gene confirmed all individuals were genetically male (n = 10) (Fig. S1C). To further investigate this, we generated *fle^DL/+^* males carrying a Y-linked RFP marker (Bernardini *et al*., 2014) and crossed homozygous *fle^DL/DL^* to *fle^DL/+^* females (Fig. Supplementary Fig. 2A, Table S1). This enabled the Y-linked RFP marker to serve as an early sex indicator, allowing sex identification of progeny from the first-instar larval (L1) stage onward. We monitored the survival of hemizygous and homozygous individuals throughout development and observed a male-to-female ratio close to 1:1 among all genotypes at the L1 larval stage (56.05 ± 2.45% and 55.53 ± 5.45% for *fle^DL/+^*and *fle^DL/DL^* males, respectively). However, only males were recovered among homozygous *fle^DL/DL^* pupae (98.65 ± 0.67%; the 1.35% female survival was hemizygous), indicating that mutant females die during larval development, likely between the L2 and L3 stages (Supplementary Fig. S2B). This finding is consistent with previous studies reporting female-specific lethality following disruption of *fle* (Krzywinska *et al*., 2021; Smidler *et al*., 2023) (Supplementary Fig. S2B, Table S1).

A phenotypic assay of the *fle* knockout strain confirmed that *fle* is a haplosufficient gene (Smidler *et al*., 2023): *fle^DL/+^* females produced larval progeny comparable to wild-type (98.7 ± 9.7 vs 83.6 ± 8.7, mean ± s.e.m., n = 40 each, p = 0.37 Dunn’s multiple comparison; Fig. 1B, Supplementary Fig. S3 and Tables S2). Similarly, *fle^DL/DL^*and *fle^DL/+^* males showed comparable larval output to wild-type, indicating no impairment of male reproductive fitness (Fig. 1B, Supplementary Fig. S3 and Tables S2).

### Generation and characterisation of the *zpg*-driven gene drive targeting *fle*

We used Recombinase-Mediated Cassette Exchange (RMCE) to replace the GFP marker in the *fle^DL^* allele with a CRISPR-Cas9 gene drive construct. The cassette comprised *cas9* under the control of the *zpg* promoter, an RFP marker (*3xP3::RFP*), and two guide RNAs (*T1* and *T14*) flanking the construct to enable multiplex targeting (Fig. 2A). We established two independent lines, *fle^zpgp#9^* and *fle^zpgp#5^*, differing only in the orientation of the genetic construct relative to the *fle* locus (Supplementary Fig. S4A-B). Hemizygous *fle^zpgp#5^* and *fle^zpgp#9^* males crossed with wild-type females exhibited no measurable reduction in fertility relative to controls (*fle^zpgp#^*^5^: 85.6 ± 6.7 versus wild-type: 93.0 ± 7.2 and *fle^zpgp#^*^9^: 106.6 ± 10.6 versus wild-type: 74.5 ± 9.0; Fig. 2B-C; Table S3-4). Both lines also displayed near-complete male transmission of the drive allele: *fle^zpgp#5^*and *fle^zpgp#9^* males transmitted the transgene to 98.0 ± 1.0% (n = 43) and 97.8 ± 0.6% (n = 32) of their progeny, respectively (mean ± s.e.m.; Fig. 2D; Table S3-4).

**Figure 2.**
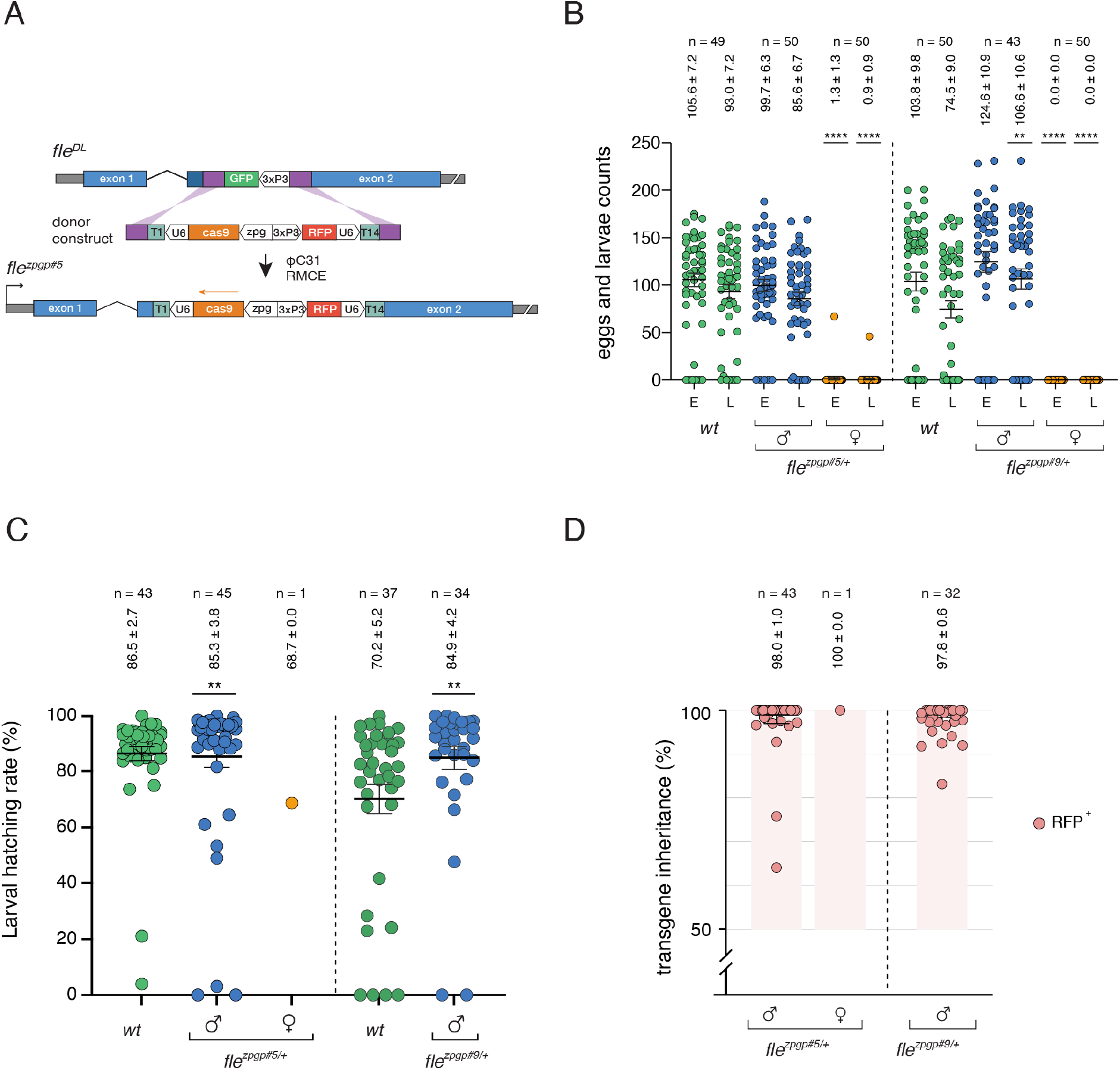
A zpg-driven gene drive targeting fle causes dominant female sterility. (**A**) Gene drive construct design and integration via ΦC31-mediated RMCE. Top: Donor construct containing *attP* sites (purple), *U6::gRNA* cassettes, *3xP3::RFP* marker, *zpg* promoter, and *cas9*. Bottom: Integrated *fle^zpgp^* allele after cassette exchange, with *Cas9* orientation indicated by the orange arrow. (**B**) Scatter plot showing the number of eggs (E) and larvae (L) produced per cross comparing *wt* control to *fle^zpgp#5/+^*(on the left) and *fle^zpgp#9/+^* (on the right) crossed to wild-type. Green, blue and orange symbols denote *wt* control, crosses with hemizygous males and crosses with hemizygous females, respectively. Egg and larvae counts were comparable between *wt* controls and hemizygous males for *fle^zpgp#5/+^* (p > 0.9999), whereas *fle^zpgp#9/+^*showed a slight increase in larvae number: p = 0.0062 (**). Crosses involving hemizygous females showed a near-complete loss of eggs and larvae compared to control (p < 0.0001, both). Kruskal-Wallis and Dunn’s multiple comparison tests. Bars show mean ± s.e.m., n = sample size. (**C**) Larval hatching rates. Hatching rates from *fle^zpgp#5/+^* and *fle^zpgp#9/+^* males crossed with wild-type do not decrease relative to wild-type. Only one *fle^zpgp#5/+^* female produced larvae, while the hatching rate of sterile *fle^zpgp#9/+^* females was not measured. Larval hatching statistics: p = 0.0031 (**) and p = 0.0019 (**) for *fle^zpgp#5/+^* and *fle^zpgp#9/+^* males, respectively, compared with wild type. Mann-Whitney (unpaired non-parametric) test. n = number of layings with at least one egg. Bars show mean ± s.e.m. (**D**) Gene drive inheritance rates based on RFP positive progeny indicate successful drive transmission via homing into the wild-type allele. *fle^zpgp#5/+^* and *fle^zpgp#9/+^* males transmit the transgene to 98.0 ± 1.0 (n = 43) and 97.8 ± 0.6 (n = 32), respectively, indicating highly efficient homing. The single fertile *fle^zpgp#5/+^* female transmitted the transgene to 100% of offspring (n = 1). Each point represents the inheritance rate from one single female lay with at least one larva. The shaded region indicates super-Mendelian inheritance. Mean ± s.e.m. values are shown.

To assess mutagenesis associated with failed homing, we analysed progeny that did not inherit the drive allele (RFP-negative offspring). Amplicon sequencing of a 476 bp region spanning both gRNA target sites was performed on RFP-negative offspring (n = 59) from crosses between *fle^zpgp#5/+^* males and wild-type females (Fig. 3A). Among 54,814 sequencing reads, 44.68% (24,493) contained modifications at one or both target sites, compared with 0.30% (169/56,818 reads) in wild-type controls (Fig. 3B; Table S5). Modified alleles were overwhelmingly out-of-frame deletions (97.72%, 23,936 reads; Fig. 3B-C). The most frequent mutation was a 79 bp deletion spanning both target sites, accounting for 34.11% of all reads, consistent with simultaneous cleavage at T1 and T14. Additional out-of-frame deletions affected either a single target site or both sites (8.82% and 1.67% of all reads, respectively; Fig. 3C). Notably, mutations localised to T1 were substantially less frequent than those at T14 (0.03% versus 8.81% of all reads; Table S5). Finally, only 5.32% of paternal alleles remained wild-type and therefore potentially susceptible to homing in subsequent generations (Fig. 3B). Overall, failed homing events predominantly resulted in loss-of-function alleles, suggesting limited opportunity for the formation of functional resistance alleles.

**Figure 3.**
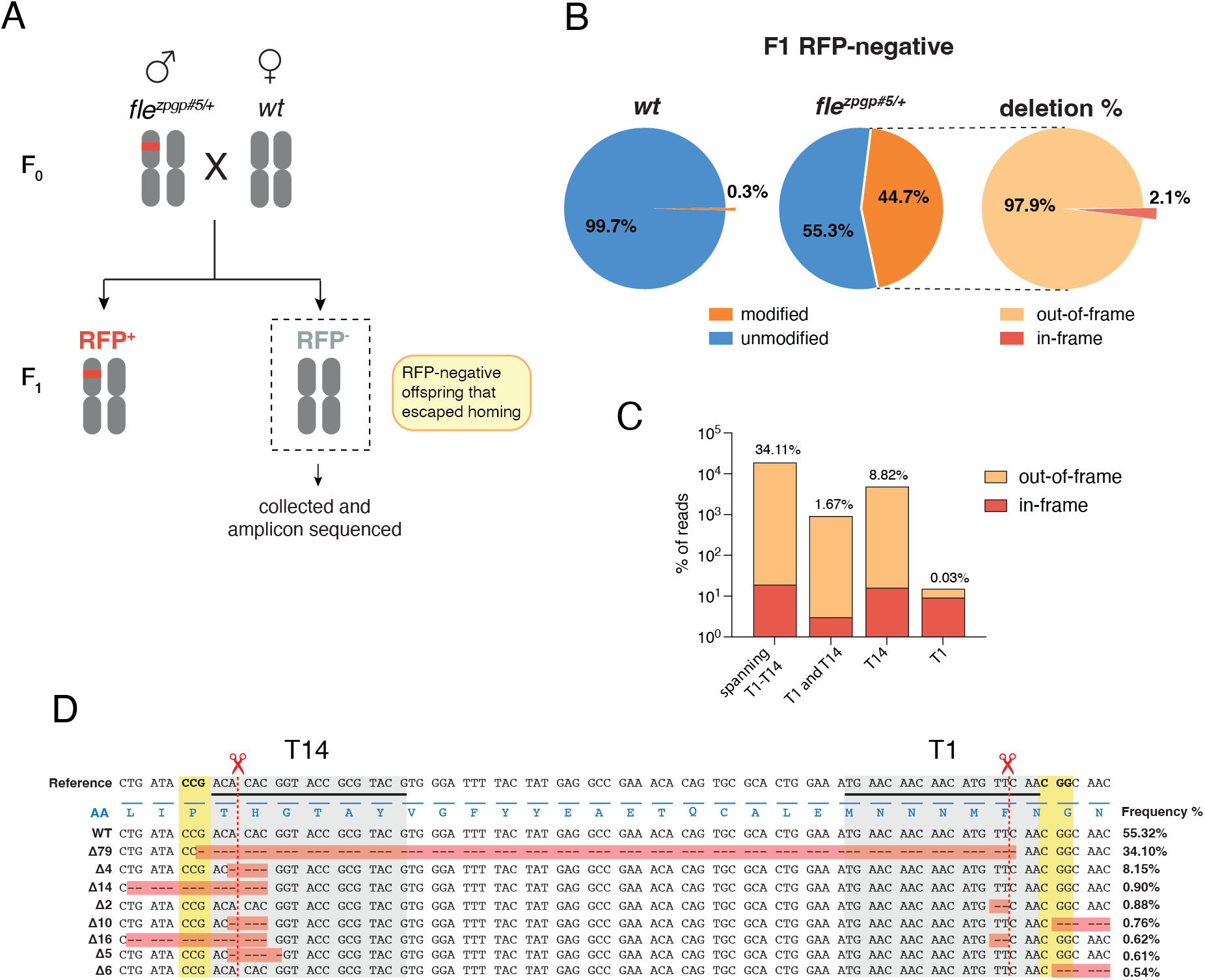
Failed homing events predominantly generate loss-of-function alleles. (**A**) Experimental design. *fle^zpgp#5/+^* males were crossed to wild-type females, and RFP-negative (inheriting the non-homing paternal allele) F1 larvae (n = 59) were collected for amplicon sequencing across both gRNA target sites. These individuals inherited a paternal chromosome that had escaped homing, allowing the characterisation of resistance-allele formation. (**B**) Mutation frequency at target sites. Pie charts show the proportion of modified versus unmodified sequencing reads. In progeny from *fle^zpgp#5/+^* fathers, 44.68% of reads carried modifications compared with 0.30% in wild-type controls (total reads = 54,814 and 56,818 for *fle^zpgp#5/+^*and wild-type, respectively). Only 5.32% of paternal alleles remained wild-type (added to the 50% of unmodified maternal reads in the pie chart), and thus susceptible to future homing. The pie chart on the right shows the percentages of in-frame (2.1%) and out-of-frame (97.9%) deletions among the modified reads. (**C**) Classification of deletion alleles. The bar graph displays the distribution of deletions by gRNA target site: T1 and T14; two independent deletions at T1 and T14 simultaneously; T14 only; and T1 only. The Y-axis uses a logarithmic scale. (**D**) Alignment of the most frequent deletion alleles to the reference *fle* coding sequence. The T14 and T1 gRNA target sites are indicated (grey shading), with PAM sites (yellow shading) and predicted cut sites (scissors, red dashed lines). The amino acid translation is shown below the reference sequence. Allele frequencies across all reads are listed to the right, including one of the 79-bp deletion predominant alleles that removes the entire region between the cut sites. Additional out-of-frame deletions affecting one (Δ4, Δ14, Δ2, and Δ5) or both (Δ10, Δ16) target sites are shown with their frequencies.

In contrast, hemizygous females of both lines were almost completely sterile. All 50 *fle^zpgp#9/+^* females failed to produce progeny, and only 1 of 50 (2%) *fle^zpgp#5/+^* laid eggs that hatched into larvae, all inheriting the transgene (n = 46) (Fig. 2B-C and Table S3-4; p<0.0001 vs wild-type, Kruskal-Wallis, Dunn’s multiple comparison test).

To investigate the basis of this sterility, the same cohort of hemizygous females that had failed to produce progeny in the fertility assays was examined for mating success. Their spermathecae were dissected five days post-mating to assess stored sperm as an indicator of successful mating (Fig. 4Ai- ii, Table S3-S4). We found that most females analysed had indeed mated, as judged by the presence of sperm in their spermathecae (*fle^zpgp#^*^5/+^: 25/38, 65.8%; Fig. 4B, Table S3 and *fle^zpgp#9/+^*: 32/44, 72.7%; Fig. 4B; Table S4). Within this mated group, sperm quantity varied, with a subset that stored reduced sperm amount (13/38 in *fle^zpgp#5/+^*, Table S3; and 19/44 in *fle^zpgp#9/+^*, Table S4; Fig. 4Ai), while the remainder retained sperm at levels comparable to control (12/38 in *fle^zpgp#^*^5/+^, Table S3 and 13/44 in *fle^zpgp#9/+^*, Table S4; Fig. 4Aii).

**Figure 4.**
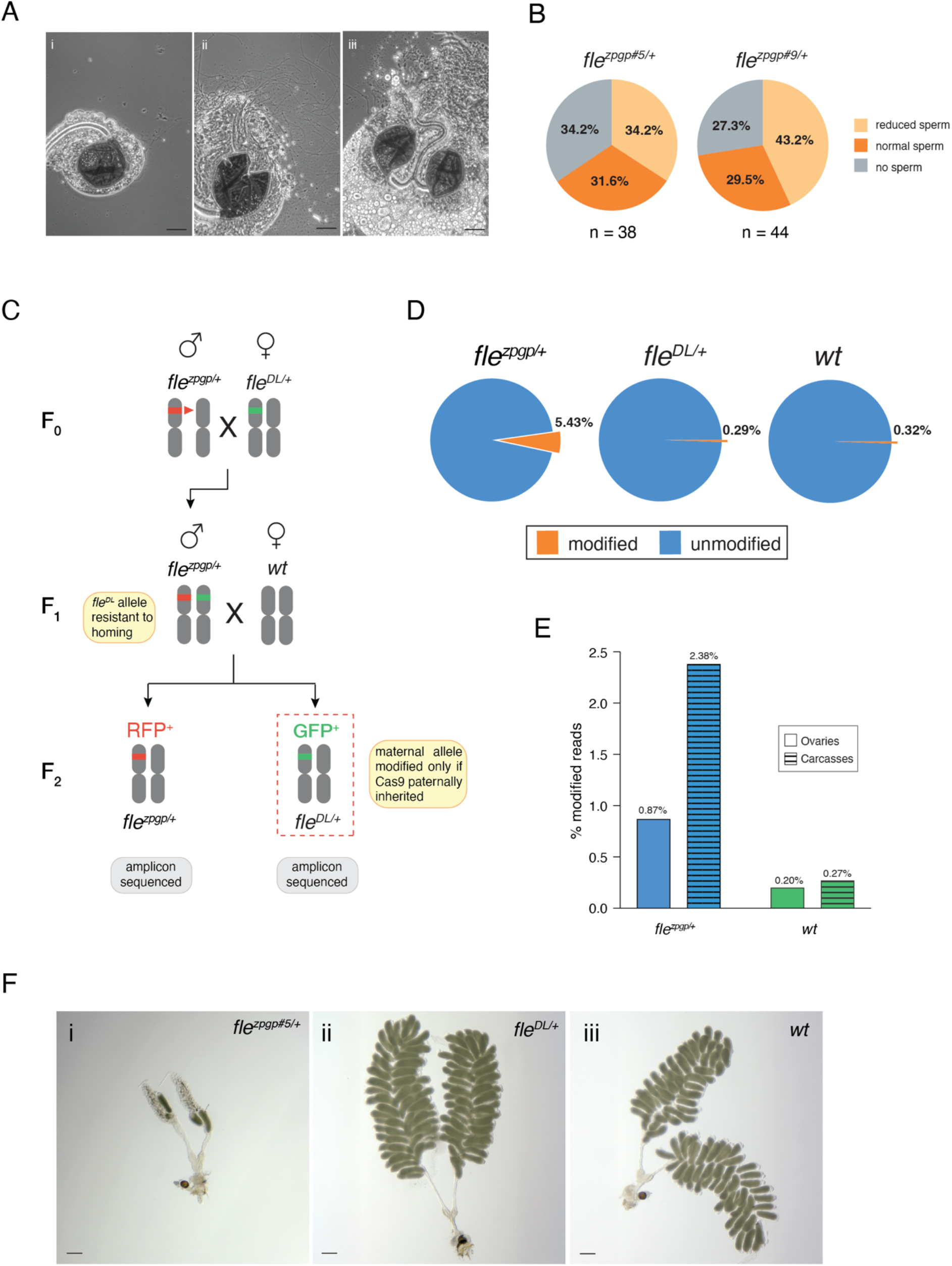
Somatic and germline Cas9 activity underlie sterility in fle^zpgp/+^ females. (**Ai-iii**) *fle^zpgp#5/+^* spermathecae analysis to assess mating status. (**i**) Spermatheca with reduced sperm content. (**iii**) Spermatheca containing abundant sperm. (**iii**) Abnormal phenotype showing two spermathecae connected to a single duct (observed in 3/34 females examined). Scale bar: 50 μm. (**B**) Pie charts showing variation in sperm quantity in the spermathecae of *fle^zpgp#5/+^* and *fle^zpgp#9/+^* females. N = number of spermathecae observed. (**C**) Experimental design to distinguish paternal Cas9 deposition from zygotic expression. *fle^zpgp/DL^* males (carrying both the gene drive and the resistant docking allele) were crossed to wild-type females. In RFP-negative offspring (which inherited *fle^DL^*but not the drive), modifications to the maternal allele can result only from paternally deposited Cas9. In RFP-positive offspring (which inherited *fle^zpgp^*), modifications indicate zygotic Cas9 activity. (**D**) Amplicon sequencing results at the pupal stage. RFP-negative (*fle^DL/+^*) offspring show modification rates (0.29%) comparable to wild-type controls (0.32%), indicating no detectable paternal Cas9 deposition. RFP-positive (*fle^zpgp/+^*) offspring show elevated modification rates (5.43%), confirming zygotic Cas9 activity. Pie charts show the proportion of modified (orange) versus unmodified (blue) reads. Data represent the mean of two biological replicates. (**E**) Tissue-specific modification rates in *fle^zpgp/+^* females. Amplicon sequencing of dissected ovaries versus carcasses (remaining body tissue) reveals higher modification in somatic tissues (2.38%) than in the germline (0.87%), demonstrating that the *zpg* promoter drives *cas9* expression in somatic tissue cells. Wild-type controls show background modification rates (0.20-0.27%). (**Fi-iii)** Overview images of dissected ovaries two days post blood meal from (**i**) *fle^zpgp#5/+^*, (**ii**) *fle^DL/+^* and (**iii**) wild-type females. Wild-type and *fle^DL/+^* ovaries contain fully developed oocytes, whereas *fle^zpgp#5/+^*ovaries are severely underdeveloped, with few mature oocytes. Scale bar 200 μm.

To further investigate and confirm that mating is not the primary cause of sterility, we conducted a further test consisting of crossing two lines of mosquitoes: one containing a transgene with the *zpg* promoter driving *cas9* (*zpg::cas9*), and the other carrying a transgene with the *T1* and *T14* gRNAs cassette (*gRNAs^fle^*), both mapping outside the *fle* locus. As with the gene drive-bearing females, trans-hemizygous *zpg::cas9/gRNAs^fle^* females crossed with wild-type showed markedly reduced fertility despite mating at a normal rate, while males were unaffected (Table S6, Supplementary Fig. S5). This independent line of evidence suggests that reduced sperm retention alone cannot fully explain the phenotype, and points to additional mechanisms acting within the female germline itself. We therefore decided to investigate the basis of this sterility more systematically.

Because both gene drive lines, *fle^zpgp#^*^5/+^ and *fle^zpgp#^*^9/+^, exhibited indistinguishable phenotypes, subsequent analyses focused on a single representative strain, *fle^zpgp#5^* (hereafter *fle^zpgp^*).

### Investigating sterility in *fle^zpgp/+^* females

The dominant sterility of *fle^zpgp/+^* females, despite the established haploinsufficiency of *fle*, suggests the involvement of at least three non-mutually exclusive mechanisms: (1) paternal deposition of Cas9 into the zygote, (2) somatic Cas9 activity generating mosaic biallelic loss-of-function mutations, and (3) germline homing that converts heterozygous to homozygous mutant cells during developmental stages in which Fle might be required for female germline development.

To investigate the basis of this sterility, we first tested whether Cas9 could be paternally deposited into the zygote, as has been reported for other gene drive systems (Hammond *et al*., 2021a; Morianou *et al*., 2024; Larrosa-Godall *et al*., 2025).

We crossed *fle^zpgp/+^* males with *fle^DL/+^* females to generate *fle^zpgp^/fle^DL^* trans-hemizygous sons carrying both the gene drive (RFP-positive) and the docking allele (GFP-positive), which is resistant to drive-mediated cleavage. These males were then crossed to wild-type females (Fig. 4C).

In GFP-positive (*fle^DL^*^/+^) offspring, the maternally inherited wild-type allele could only be mutagenised by Cas9 deposited from the father (Fig. 4C). Amplicon sequencing of the target site revealed very low mutation rates in GFP-positive progeny, comparable to background levels (*fle^DL/+^*, 0.29% vs 0.32% in wild-type) (Fig 4D), effectively ruling out paternal deposition of Cas9. In contrast, RFP-positive progeny (*fle^zpgp/+^*) showed significantly elevated mutation rates at the target site (5.43 ± 0.14%, n = 2 replicates), suggesting potential leaky *cas9* expression in individuals carrying the gene drive allele (Fig 4D).

To determine whether Cas9 activity in *fle^zpgp/+^* extended to somatic tissues and was therefore not restricted to germline tissues, we compared the rate of NHEJ modifications between dissected ovaries and the remaining body parts (carcasses) of *fle^zpgp/+^* females, using wild-type females as controls. Amplicon sequencing of the *fle* target sites in the two samples showed that somatic tissues had marginally higher modification rates (2.38%, n = 2 technical replicates, 20 individuals) than ovarian tissues (0.87%, n = 2 technical replicates, 20 pairs of ovaries; Fig 4E), indicating that the *zpg* promoter, although described as germline-specific, exhibits some activity in somatic tissue. To assess whether germline homing could account for the sterility observed in *fle^zpgp/+^* females, we examined the ovaries of blood-fed individuals for developmental disruption that might result from conversion of heterozygous cells to homozygous mutant cells during stages when *fle* might be critical for germline development. All *fle^zpgp/+^* ovaries examined (n = 38; Table S3) were underdeveloped, with few or no mature oocytes (Fig. 4F*i-iii*). Occasionally, we observed ovaries with two spermathecae (3/34 females; Fig. 4Aiii). Confocal microscopy revealed follicles with excess nurse cells (Supplementary Fig. S6B and E) and increased stalk cells connecting germaria to secondary follicles (Supplementary Fig. S6C and F). Together, these ovarian defects are consistent with biallelic loss of *fle* function in the germline cells, supporting germline homing as a major cause of sterility in *fle^zpgp/+^* females.

### Delaying Cas9 expression with the *spo11* promoter partially restores female fertility

Having established that germline homing is a major driver of sterility in *fle^zpgp/+^* females, we asked whether modifying the timing of *cas9* expression could mitigate this effect.

Although stage-specific expression data during oogenesis are not currently available for *An. gambiae*, single-cell RNA sequencing data from male gametogenesis indicate that *fle* is expressed during the primary spermatogonial stages (Page *et al*., 2023), when *zpg* is active. If a similar expression pattern occurs during female germline development, the homing-mediated loss of Fle function during early oogenesis could explain the sterility phenotype observed in gene drive females.

We hypothesised that restricting *cas9* expression to stages following early germ cell specification would preserve female fertility while retaining high homing efficiency.

To test this, we designed a gene drive construct where we replaced the *zpg* promoter with sequences from the *sporulation 11* (*spo11*) gene, which is essential for meiotic recombination (Atcheson *et al*., 1987; Dernburg *et al*., 1998; McKim and Hayashi-Hagihara, 1998; Hartung *et al*., 2007). The *spo11* promoter is active later in germline development than *zpg* and shows elevated expression in females after blood feeding (Taxiarchi *et al*., 2019; Terradas *et al*., 2021; Page *et al*., 2023). Recently, a drive system using *spo11::cas9* to target *dsx* (*gRNA^dsx^*) demonstrated that this promoter can support efficient homing in the germline of both sexes; however, the transmission rates achieved were variable, depending on the sex of the carrier and the construct’s genomic location (Grilli *et al*., 2026). We generated the *fle^spo11p^* construct by replacing the *zpg* promoter and terminator with *spo11* regulatory sequences while maintaining identical *cas9*, gRNA cassettes (*T14* and *T1*), and RFP marker sequences (Fig. 5A). Multiple independent lines were obtained through RMCE, of which we retained two representatives of the alternative *cas9* orientations, *fle^spo11p#3^* and *fle^spo11p#6^* (Supplementary Fig. S7A-B). These mosquito lines showed different male transmission rates in initial bulk test crosses (97.5% and 77.5%, respectively; Supplementary Fig. S7C). For detailed characterisation, we focused on the *fle^spo11#3^*line (hereafter *fle^spo11p^*), which showed a higher male transmission rate (Supplementary Fig. S7C).

**Figure 5.**
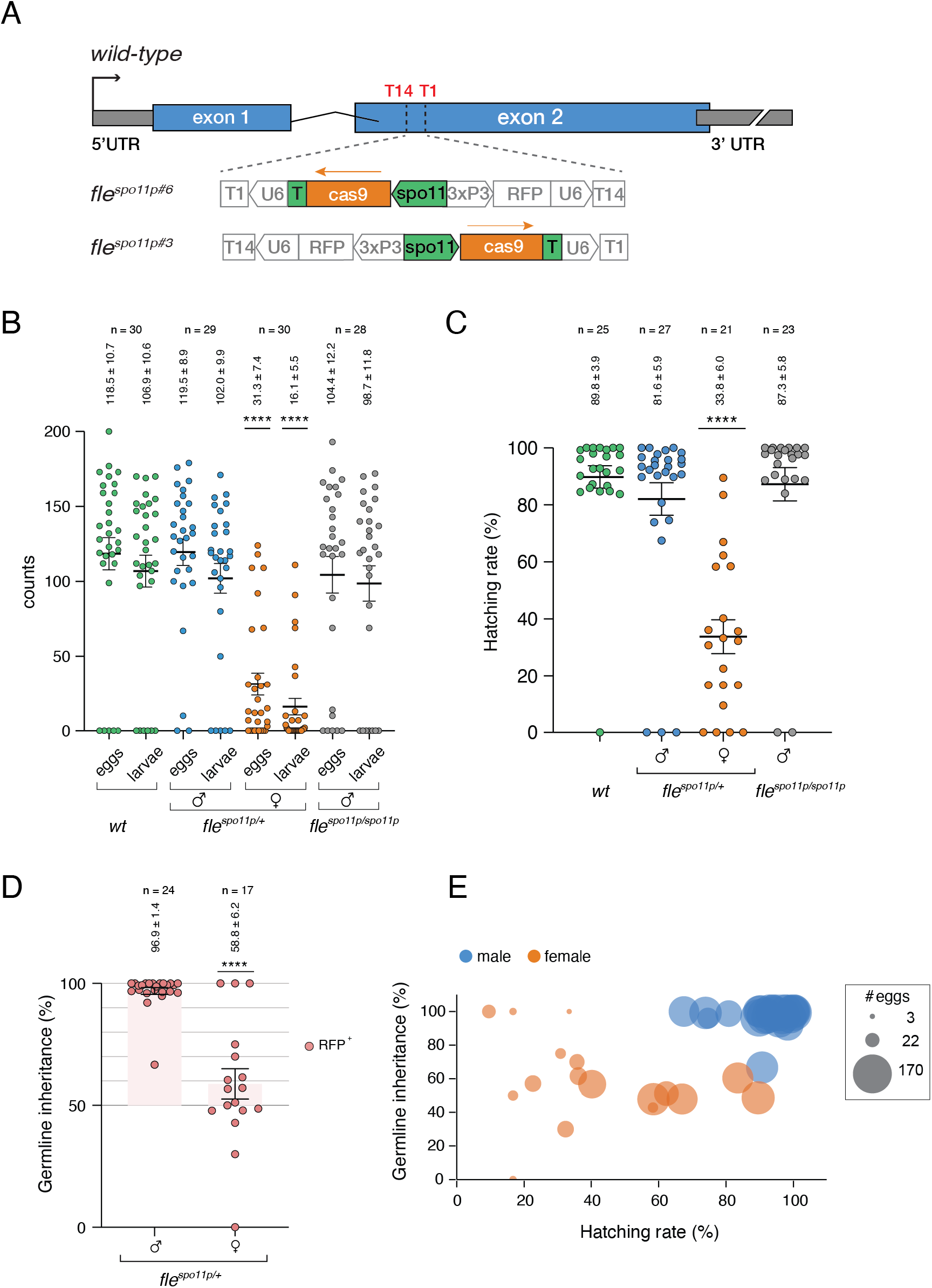
The spo11 promoter partially restores female fertility. (**A**) *fle^spo11p^* gene drive constructs showing the orientations of *cas9* (green) in the *fle^spo11p#3^* and *fle^spo11p#6^* lines. The constructs share all components with the *fle^zpgp^* transgene: *U6::gRNA* cassettes and the *3xP3::RFP* marker, except for the spo11 promoter and terminator and are integrated at the same *fle* locus position via ΦC31-mediated RMCE. (**B**) Fertility of *fle^spo11p^* individuals. Wild-type females mated to *fle^spo11p/+^* males show normal eggs (119.5 ± 8.9) and larval (102.0 ± 9.9) output. *fle^spo11p/+^* females crossed to wild-type males show significantly reduced but detectable fertility (eggs: 31.3 ± 7.4; larvae: 16.1 ± 5.5), in contrast to the complete sterility observed with *fle^zpgp^*. Each point represents the progeny from a single female; p < 0.0001 (Kruskal-Wallis test with Dunn’s correction versus wild-type. (**C**) Larval hatching rate from *fle^spo11p/+^* males (81.6± 5.9%) is comparable to wild-type (89.8 ± 3.9%). Hatching from *fle^spo11p/+^* females is reduced (33.8 ± 6.0%), consistent with partial fertility. p < 0.0001 versus wild-type. (**D**) Gene drive inheritance rates. *fle^spo11p/+^* males transmit the transgene to 96.9 ± 1.4% of offspring (n = 24), comparable to *fle^zpgp^*. However, females transmit to only 58.8 ± 6.2 of their offspring (n = 17), near Mendelian values, indicating minimal homing through females. Each point represents the percentage of gene drive alleles in a single laying; the dashed background indicates > 50% Mendelian inheritance. ****p < 0.0001 (Mann-Whitney test). Bars show mean ± s.e.m. (**E**) Bubble plot showing germline inheritance (%) as a function of hatching rate (%) for *fle^spo11p/+^*females (orange bubbles) and males (blue bubbles) crossed to wild-type partners. Bubble size is proportional to the number of eggs deposited per individual cross. Female drive carriers exhibit highly variable germline inheritance and hatching rates, with extreme values (0% and 100%) predominantly associated with crosses that yield very few eggs (small bubbles) and low inheritance rates. Male drive carriers show consistently high germline inheritance (93-100%) across uniformly high hatching rates (60-100%) with larger clutch sizes.

To assess the fitness consequences of delayed *cas9* expression, we generated heterozygous and homozygous carriers of the *fle^spo11p^* gene drive and performed fertility assays. Both hemizygous and homozygous *fle^spo11p^* males exhibited fertility levels comparable to those of wild-types, with no significant differences in egg production or hatching rates (Kruskal-Wallis and Dunn’s multiple comparison tests, p > 0.9999) (Fig. 5B-C, Table S7).

In contrast, hemizygous *fle^spo11p^* females exhibited a partial rescue of fertility relative to the complete sterility observed in hemizygous *fle^zpgp^* females. However, both egg production and hatching rates remained substantially reduced compared with wild-type females (31.3 ± 7.4 eggs and 33.8 ± 6.0 hatching rate versus 118.5 ± 10.7 eggs and 89.8 ± 3.9 hatching rate, respectively, mean ± s.e.m., n = 30; Kruskal-Wallis and Dunn’s multiple comparison tests, p < 0.0001 for both comparisons) (Fig. 5B-C). These findings support the hypothesis that the dominant sterility phenotype associated with the *zpg* system arises from loss of *fle* function in the female germline and suggest an essential role for *fle* in female reproductive development beyond its established function in sex determination.

We could not analyse homozygous *fle^spo11p^* females because they died at the early larval stage, phenocopying *fle^DL/DL^* homozygotes and confirming that complete loss of *fle* function is lethal. Male transmission of the *fle^spo11p^*construct remained high (96.9 ± 1.4%, mean ± s.e.m., n = 24), comparable to that of the *zpg*-driven system, indicating efficient homing in the male germline (Fig. 5D and E). By contrast, transmission through *fle^spo11p/+^* females remained close to Mendelian rates (58.8 ± 6.2%, *n* = 17) with only a few females showing high inheritance rate but also a markedly reduced number of eggs and hatching rates (Fig. 5D-E).

To further characterise Cas9 activity at the target sites, we examined the mutation spectrum in RFP-negative individuals that did not inherit the drive. In the offspring of *fle^spo11p/+^* males crossed to wild-type, mutations were detected at a rate of 15.08% (n = 40 individuals) compared to 0.30% in wild-type (Fig. 6A and Table S8). Among these, out-of-frame mutations were considerably more prevalent than in-frame mutations (97.05%, 14392 reads vs 2.95%, 437 reads, respectively; Table S8, Fig 6B), a pattern similar to the *zpg* system, where modifications are also predominantly out-of-frame (99.7% of modified alleles).

**Figure 6.**
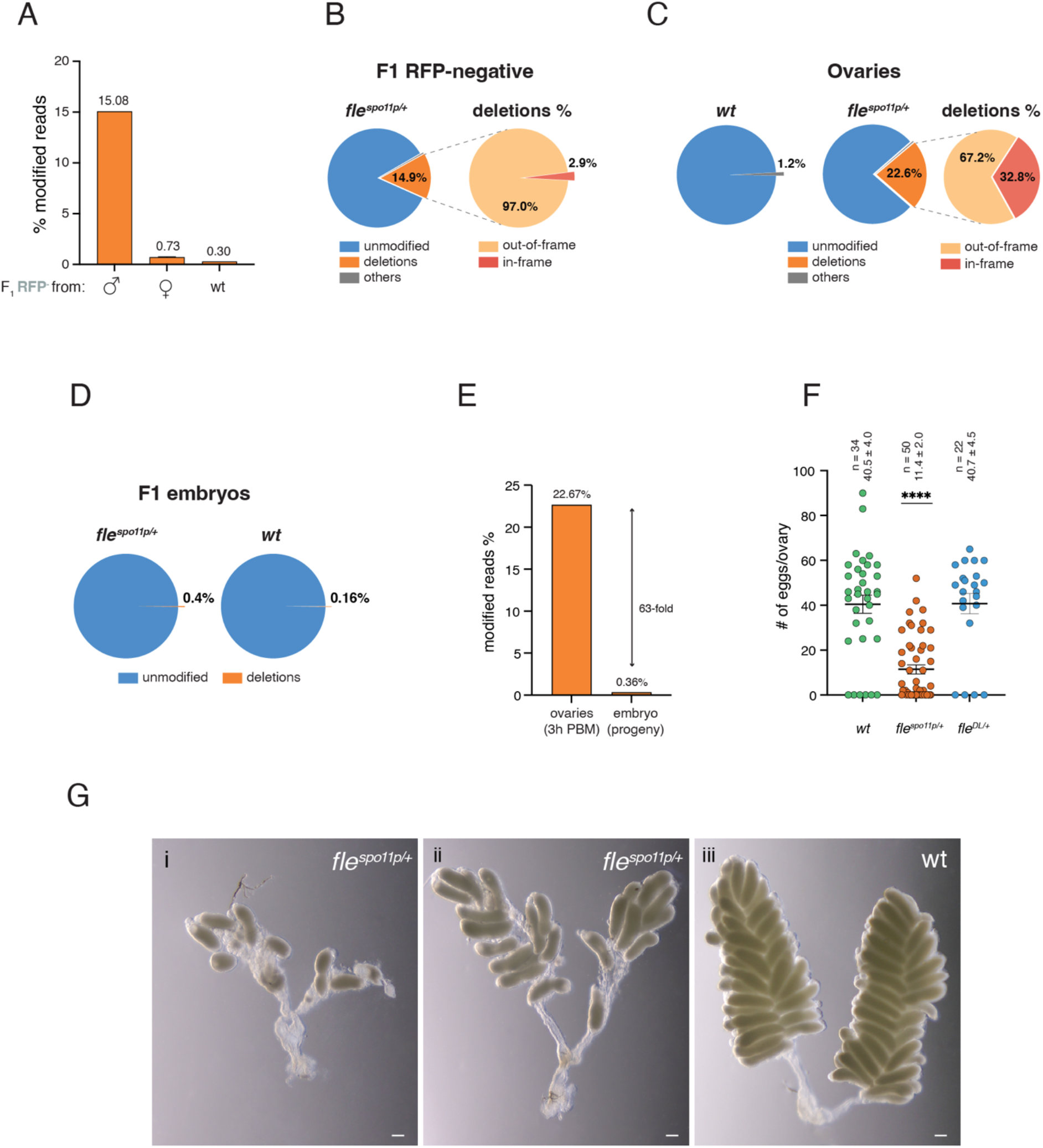
Oocytes carrying mutated fle alleles are eliminated before oviposition. (**A**) Bar graph showing the percentages of modified alleles in RFP-negative offspring from *fle^spo11p/+^*parents and wild-type. Male parents transmit modified alleles to 15.08% of their offspring (pooled across 40 individuals). In contrast, female parents transmit modified alleles to only 0.73% of offspring (pooled across 110 individuals), comparable to the 0.30% background in wild-type controls. This indicates that oocytes carrying *fle* mutations are eliminated or fail to develop. (**B**) Editing outcomes in F1 RFP-negative offspring from *fle^spo11p/+^*fathers. Pie charts showing the distribution of allele classes, with deletion comprising 14.9% of total alleles and a negligible fraction classified as “others” (SNPs and substitutions). The deletion fraction is expanded to show that the majority are out-of-frame mutations (97.0%) with a minor fraction of in-frame events (2.9%). Colours denote unmodified (blue), deletions (orange), other alleles (grey), out-of-frame (light orange) and in-frame (red). Dashed lines indicate that the right panel expands on the deletion section from the left panel. (**C**) Editing outcomes in the ovaries of wild-type and *fle^spo11p/+^* individuals (n = 20 each; 3 hours post-blood meal). Pie charts show the distributions of sequence outcomes following genome editing. In wild-type ovaries (*wt*), most alleles remain unmodified, with a small fraction classified as “others” events (1.2%, SNPs and substitutions). In *fle^spo11p/+^* ovaries, deletions constitute 22.6% of alleles, with the remainder unmodified except for 0.1% classified as “others”. The deletion category (highlighted and expanded) is further divided into out-of-frame (67.2%) and in-frame (32.8%) events. Colours denote unmodified (blue), deletions (orange), other alleles (grey), out-of-frame (light orange) and in-frame (red). Dashed lines indicate that the right panel expands on the deletion section from the left panel. (**D**) Editing outcomes in F1 embryos from *fle^spo11p/+^* and wild-type mothers. Pie charts showing allele distributions in 24-hour-old embryos derived from *fle^spo11p/+^* (left) and wild-type (right). In both cases, the vast majority of alleles are unmodified (blue) with rare deletions (orange). The frequency of deletions is modestly increased in embryos from *fle^spo11p/+^* mothers (0.4%) compared with that in wild-type embryos (0.16%). (**E**) Enrichment of modified reads in ovaries compared to progeny embryos. Bar plot showing the percentage of modified alleles detected in ovaries at 3 hours post blood meal (PBM) and in embryos (progeny). Ovaries exhibit a markedly higher fraction of modified reads (22.67%) than embryos (0.36), corresponding to approximately 63% enrichment (χ² = 27511, df =1, p > 0.0001). Values above bars indicate percentages and the double-headed arrow denotes the fold difference between conditions. (**F**) Loss of *fle* function reduces oogenesis output. Dot plots showing the number of eggs per ovary (48 hours post blood meal) in three genotypes: wild-type, *fle^spo11p/+^* and *fle^DL/+^*. Each dot represents an individual ovary. Sample sizes (n) and mean ± s.e.m. values are indicated above each group. The *fle^spo11p/+^* genotype exhibits a marked reduction in egg number compared with both wild-type and *fle^DL/+^*controls. Statistical significance was assessed using the Mann-Whitney U test (****p<0.0001). (**G**) Ovarian phenotypes in *fle^spo11p/+^*females. Ovaries two days post blood meal from *fle^spo11p/+^* females (i and ii) and from wild-type (iii). *fle^spo11p/+^*ovaries show underdeveloped follicles and few mature oocytes, consistent with the elimination of those that successfully homed, generating homozygous mutant cells.

In contrast, the rate of mutations in RFP-negative offspring from *fle^spo11p/+^* females was only 0.73% (n = 110 RFP-negatives), a frequency comparable to the background mutation rate observed in wild-type controls (0.34%, *n* > 50; Fig. 6A). An analogous pattern was observed in the independent *fle^spo11p#6^* line (Supplementary Fig. S8A). Together, these results indicate that the *spo11* promoter supports efficient Cas9 activity in the male germline but only limited activity in females (Grilli *et al*., 2026).

### Oocytes carrying mutated *fle* alleles are eliminated before oviposition

Although the marked difference in drive transmission between males and females suggested reduced activity of the *spo11*-driven construct in the female germline, we tested an alternative hypothesis: that homing occurs efficiently in female germ cells, but oocytes carrying biallelic loss-of-function *fle* alleles are subsequently eliminated during oogenesis, as proposed to explain the complete sterility observed in the *zpg*-driven system. In this scenario, efficient homing would not increase transmission, since mutant germ cells would fail to contribute to the next generation. The partial restoration of fertility observed in *fle^spo11p/+^* females could then be attributed to the later onset of *spo11*-driven *cas9* expression, which might allow some germ cells with a functional *fle* allele to complete development and contribute to reproduction.

To test this hypothesis, we performed amplicon sequencing of the *fle* target site in the ovaries of *fle^spo11p/+^* individuals. Although this assay cannot quantify the extent of homing, it can detect mutations indicative of Cas9 activity. We reasoned that mutations should be detectable in ovarian tissue but largely absent from viable offspring if homing occurred and mutant oocytes were selectively eliminated.

Ovaries from mated *fle^spo11p^*^/+^ females were dissected three hours post-blood meal, a time point when *spo11* is active and drives *cas9* expression (Terradas *et al*., 2021). Target-site deletions were detected in 22.56% of reads, more than 18-fold higher than the 1.21% modifications observed in wild-type ovaries. Frameshifts were twice as frequent as in-frame deletions (67.2% and 32.8%, respectively; Fig. 6C and Table S9), providing direct evidence that Cas9-mediated cleavage and mutagenesis occur in the ovarian germline.

By contrast, analysis of 24-h-old F1 embryos from *fle^spo11p/+^* females crossed with wild-type males showed only 397 modified reads, corresponding to 0.36 ± 0.18% of all reads (n = 2 embryo-batch replicates), comparable to wild-type embryos (0.16%, n = 1 embryo batch; Fig. 6D; Table S10) and representing a > 60-fold reduction relative to ovaries (Chi-square 27511, df = 1, p < 0.0001; Fig. 6E). The near absence of mutant alleles in embryos indicates that selection occurs before oviposition rather than through post-fertilisation embryonic lethality. Consistent with this model, ovaries dissected from *fle^spo11p/+^* females at 3 days post-blood meal contained significantly fewer oocytes than wild-type ovaries (11.4 ± 2.0, n = 50 versus 40.5 ± 4.0, n = 34, respectively; Mann–Whitney two-tailed test, P < 0.0001; Fig. 6F-G), indicating substantial loss of developing oocytes following disruption of *fle*.

### Cas9 inhibition by AcrIIA4 confirms its role in *fle*-mediated sterility

Together, these results indicate that *fle* disruption in the female germline triggers the elimination of germ cells during development. Using two independent germline promoters active at different stages of germline differentiation, we established that Cas9-mediated mutagenesis of *fle* underlies the observed sterility in females. To further validate this conclusion, we employed AcrIIA4, an anti-CRISPR protein that specifically inhibits Cas9 activity (Fig 7A). We generated *fle^zpgp^*; *vasa2::AcrIIA4* individuals by crossing hemizygous *fle^zpgp^* males with females carrying a transgene in which the *vasa2* promoter drives *AcrIIA4* expression (Taxiarchi *et al*., 2021) (Fig. 7A). Because *vasa2* is expressed in the germline earlier than *zpg* and also partially overlaps with it, AcrIIA4 under *vasa2* control was expected to inhibit Cas9 as soon as the protein became available, providing a stringent test of whether AcrIIA4 activity is sufficient to restore fertility. We then compared the fertility of trans-hemizygous *fle^zpgp^*; *vasa2::AcrIIA4* males and females with that of hemizygous *fle^zpgp^*males and females. Trans-hemizygous *fle^zpgp^*; *vasa2::AcrIIA4* females exhibited restored fertility, producing an average of 36.4 ± 8.8 eggs with a larval hatching rate of 58.9 ± 8.8% (Fig. 7B; Table S11). In contrast, hemizygous *fle^zpgp^* females laid no eggs (Fig. 7B-C; Table S11). Moreover, no homing was detected in either trans-hemizygous males or females, with mean inheritance rates of 48.2 ± 1.6 and 49.9 ± 5.7, respectively, compared with 100% in hemizygous *fle^zpgp^* males, confirming the Cas9-inhibitory role of AcrIIA4 (Fig 7D; Table S11).

**Figure 7.**
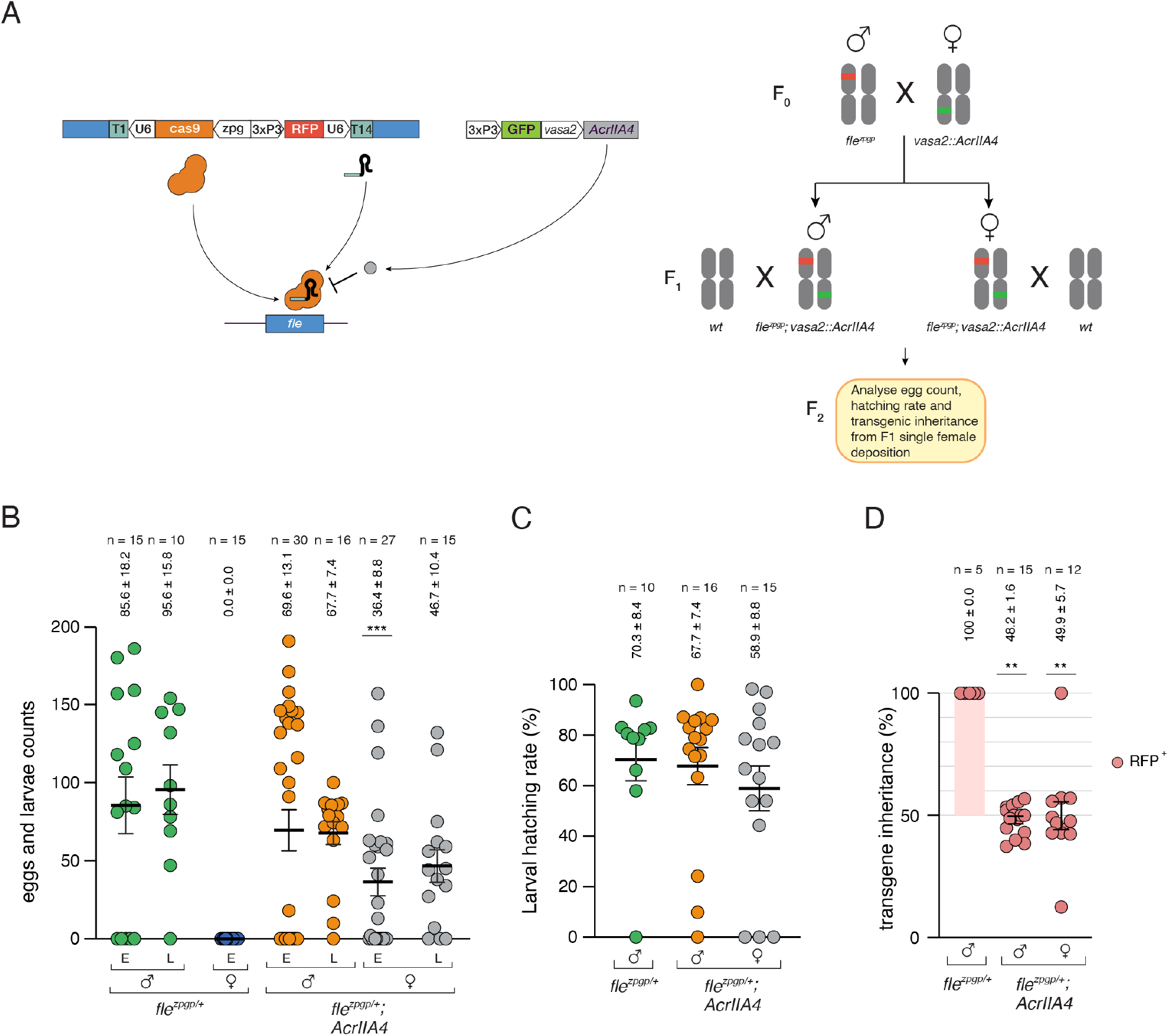
AcrIIA4-mediated Cas9 inhibition rescues female fertility in fle^zpgp/+^ individuals. (**A**) Schematic of AcrIIA4 inhibition of the *fle*-targeting Cas9-gRNA ribonucleoprotein complex. Cas9 protein (orange) loaded with the specific guide RNA (teal) binds and cleaves the target site within the endogenous *fle* locus (blue bar). AcrIIA4 (grey circle), expressed from the *vasa2::AcrIIA4* transgene, blocks the association of the Cas9-gRNA complex with target DNA preventing cleavage of *fle.* (**B**) Crossing scheme used to test the effect of AcrIIA4 on drive transmission. F_0_ males hemizygous for the *fle^zpgp^* drive allele (RFP chromosome mark) were crossed to F_0_ females hemizygous for *vasa2::AcrIIA4* (GFP chromosome mark). Trans-hemizygous *fle^zpgp^; vasa2::AcrIIA4* F_1_ males and females were each backcrossed to wild-type (*wt*) partners and the F2 progeny from single females was scored for egg number, larvae hatched and transgene inheritance (RFP-positives) to determine whether AcrIIA4 co-expression suppresses super-Mendelian transmission of the *fle* drive allele. (**C**) Eggs and larval progeny from individual layings across four cross types (left to right): *fle^zpgp/+^* and *fle^zpgp/+^; vasa2::AcrIIA4/+* males and females crossed to wild-type partners. *fle^zpgp/+^*females are completely sterile (E = 0.0 ± 0.0), whereas *fle^zpgp^; vasa2::AcrIIA4* females produce both eggs and larvae (E = 36.4 ± 8.8 and L = 46.7 ± 10.4), indicating that Cas9 inhibition restores fertility. Male fecundity is comparable across genotypes. Each dot represents an individual laying (n = number of laying). Horizontal lines indicate mean ± s.e.m., with values shown above each group. Eggs and larvae output was compared between *fle^zpgp/+^*and *fle^zpgp/+^; vasa2::AcrIIA4/+* males (P = 0.5643 and 0.1779, respectively) and between *fle^zpgp/+^* and *fle^zpgp/+^; vasa2::AcrIIA4/+* female eggs (***P = 0.0005)

### Population suppression modelling predicts feasible release requirements for *fle^zpgp^*

The combination of complete female sterility in *fle^zpgp/+^* mosquitoes and near-complete transmission through males (98%) suggests that the system could function as a male-driven population-suppression strategy (Labbé *et al*., 2012; Chen *et al*., 2024; Strampelli *et al*., 2025; Tolosana *et al*., 2025). To evaluate its potential impact at the population level, we used computational modelling to estimate the release rates required to achieve substantial suppression.

The model predicts that repeated releases of heterozygous *fle^zpgp/+^* males at 16.9% of the wild-type male population per generation would reduce the female biting population by 95% within 36 generations (Fig. 8 and Table S12). This release threshold is substantially higher than that predicted for an idealised male-drive female-sterile (MDFS) system targeting *fle*, which assumes perfect homing and the absence of resistant alleles and would require releases of only 4.3%.

**Figure 8.**
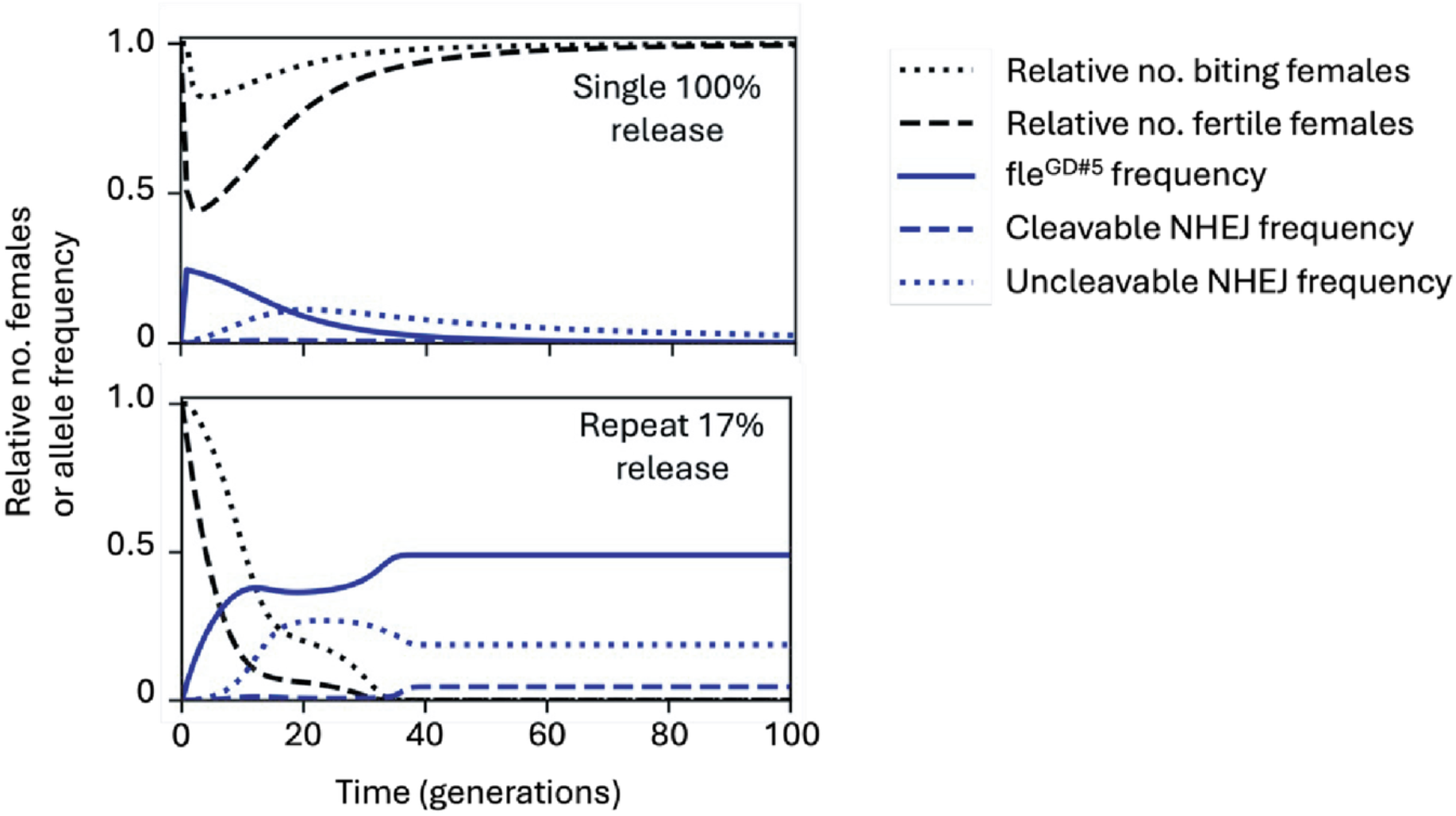
Population suppression modelling shows repeated releases of fle^zpgp^ males can achieve sustained suppression. Population dynamics modelling comparing single- and repeated-release strategies. Lines show relative abundance of fertile females (black, dashed), biting females (black, dotted), *fle^zpgp^*drive allele frequency (blue solid), cleavable NHEJ products (at least one unmodified gRNA, blue dashed), and uncleavable NHEJ products (both gRNA modified, blue dotted) over 100 generations. **Top panel (single 100% release):** A single large release of hemizygous at 100% of initial male population size produces only transient suppression. The drive allele frequency increases to ∼25%, but declines as resistance alleles accumulate through NHEJ. Both fertile and biting female populations recover to pre-release levels after approximately 50 generations, demonstrating that single releases cannot achieve sustained suppression even at very high introduction rates. **Bottom panel (Repeated 17% releases):** Sustained releases of hemizygous at 17% of the initial male population size per generation for 36 generations achieve effective population suppression. The drive allele frequency stabilises at ∼47%, maintaining the number of fertile females near zero and reducing the biting female population by >95%. Despite the ongoing accumulation of resistance alleles, the repeated introduction of drive alleles maintains sufficient genetic load to suppress reproduction. Drive inheritance and fitness parameters are based on empirical observations, and the population is assumed to have an R_m_ of 6.

Model analysis identified alleles resistant to cleavage at both target sites (A4) as the principal factor limiting suppression efficiency. Because these alleles are refractory to further cleavage, they block homing in males and reduce drive transmission. Consistent with this interpretation, the required release rate decreased to 4.5% when all NHEJ-derived alleles were assumed to remain susceptible to cleavage (p = 1), approaching the performance of the idealised MDFS system.

## Discussion

By comparing homing gene drives in which *cas9* expression was controlled by germline promoters with distinct temporal activity, we uncovered a previously unrecognised role for *femaleless* (*fle*) during female germline development. Although *fle* has previously been characterised as a master regulator of female sex determination in *Anopheles gambiae*, our results demonstrate that it is also essential for oogenesis. Early *cas9* expression driven by the *zpg* promoter resulted in complete sterility in hemizygous females, whereas delaying cleavage with the later-acting *spo11* promoter partially restored fertility. Together, these findings indicate that the developmental timing of *fle* disruption, rather than loss of function alone, determines female reproductive outcome and reveal an essential requirement for *fle* during female germline development.

The contrasting phenotypes produced by the two promoters provide insight into the developmental window during which Fle is required. The *zpg* promoter is active early in germline development and, in males, its expression overlaps with *fle* during the primary spermatogonial stages (Terradas *et al*., 2021; Page *et al*., 2023). If a similar temporal relationship exists during oogenesis, Cas9-mediated cleavage would occur during germline stem-cell divisions or early follicle development, before oocyte differentiation. Homing and error-prone repair at this stage would generate biallelic loss-of-function mutations within individual germ cells, disrupting *fle* before oocyte development is established. Consistent with this interpretation, ovaries from *fle^zpgp/+^* females were uniformly abnormal, displaying severe developmental defects characterised by the absence of mature oocytes, abnormal follicles containing excess nurse cells, and increased numbers of stalk cells connecting germaria to secondary follicles. These defects strongly suggest that sterility results from disruption of *fle* within the germline itself rather than from defects in mating or adult female physiology.

In contrast, delaying Cas9 expression using the *spo11* promoter allowed early germline development to proceed in the presence of a functional *fle* allele before mutagenesis occurred. Although fertility was only partially restored, this temporal shift was sufficient to demonstrate that *fle* function is specifically required during early oogenesis. The apparent paradox of efficient Cas9 activity in males but near-Mendelian transmission of the drive through females was resolved by amplicon sequencing of the *fle* target site. While modified alleles accumulated at high frequency in ovarian tissue during the peak of *spo11* activity, they were almost absent from embryos, indicating that mutant germline cells are eliminated before oviposition rather than through post-fertilisation embryonic lethality. The marked reduction in mature oocytes in *fle^spo11p/+^* females further supports this model (Fig. 9).

**Figure 9.**
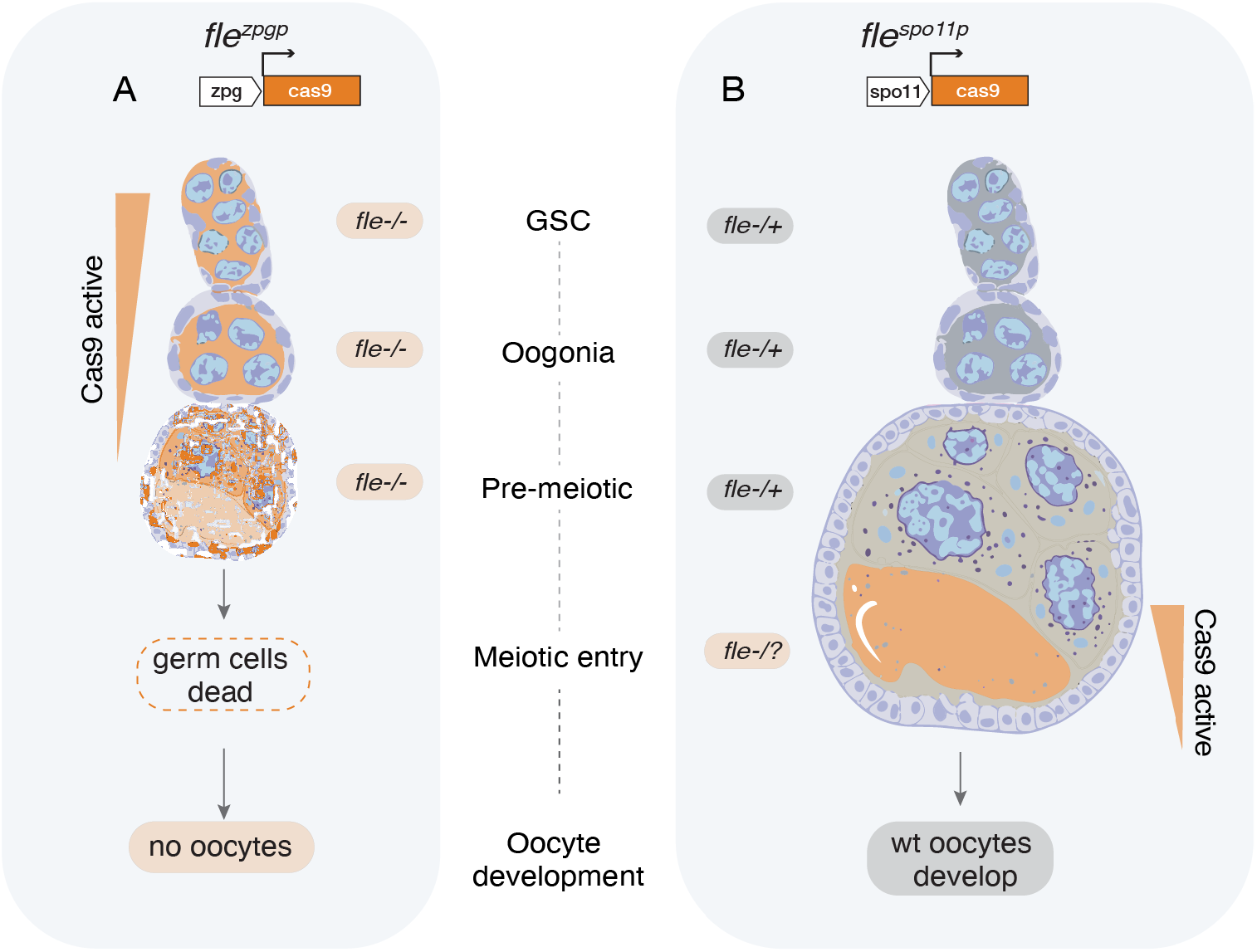
fle disruption outcomes in fle^zpgp/+^ and fle^spo11p/+^ during egg development. Expression patterns of *zpg*- and *spo11*-*cas9* in *A. gambiae* ovariole following a blood meal, adapted from Terradas *et al*. (Terradas *et al*., 2021). (**A**) In the *fle^zpgp^*construct, the *zpg* promoter initiates *cas9* expression from the earliest germline stem cell stages (GSC). Biallelic disruption of *fle* (*fle*^-/-^) occurs pre-meiotically, preventing germ cell differentiation into oocytes. (B) In the *fle^spo11p^* construct, the *spo11* promoter induces *cas9* expression at the onset of meiosis. GSCs, oogonia, and pre-meiotic germ cells retain a functional *fle* allele (*fle^-/+^*), permitting normal early oogenesis. Upon Cas9 activation after the blood meal at meiotic entry, three outcomes are possible depending on the fate of the second *fle* allele (*fle^-/?^*): successful gene drive homing, NHEJ mutation disrupting *fle* function or unsuccessful homing that preserves *fle* function and supports oocyte development. Consequently, oocytes with functional *fle* alleles can be fertilised and produce progeny. Orange shading represents Cas9 expression. Vertical orange wedges indicate the developmental window of Cas9 activity for each promoter.

Our findings also reiterate an important principle for homing-based gene-drive target selection. Candidate population suppression genes are commonly prioritised according to whether they produce strong female-specific phenotypes while remaining haplosufficient at the organismal level. However, our results demonstrate that this criterion alone is insufficient. Although *fle* is haplosufficient for female development, homing converts heterozygous germ cells into homozygous mutant ones. If the targeted gene has a cell-autonomous role during gametogenesis, these mutant germ cells are selectively lost, preventing efficient transmission through females. Thus, genes that appear suitable targets based on organismal phenotypes may nevertheless impose an intrinsic barrier to the spread of self-sustaining homing gene drives.

The dual role of *fle* in female sex determination and dosage compensation makes it functionally analogous to *Sex lethal* (*Sxl*) in *Drosophila melanogaster*, despite the absence of direct orthology (Steinmann-Zwicky, 1994; Hashiyama *et al*., 2011; Rose *et al*., 2020; Krzywinska *et al*., 2021; Scott, 2021; Grmai *et al*., 2023). Notably, targeted disruption of *Sxl* in the *Drosophila* germline similarly causes ovarian tumours, defective cyst differentiation and female sterility (Salz, 1992; Goyal et al., 2024). The similarities between these phenotypes and those observed after *fle* disruption suggest that the intimate relationship between sex-determination pathways and germline developmental competence may represent a conserved feature of insect reproductive biology rather than a lineage-specific property of mosquitoes. This essential role of *fle* in *Anopheles gambiae* oogenesis may therefore reflect a conserved mechanistic link between the sex-determination regulatory hierarchy and germ cell developmental competence.

From an applied perspective, the requirement for *fle* during oogenesis fundamentally alters the behaviour of a homing suppression drive. Rather than functioning as a self-sustaining gene drive, the *fle^zpgp^* construct behaves as a male-driven, female-sterile release system analogous to the Male Drive Female Sterile (MDFS) strategy previously developed by targeting *doublesex* (Strampelli *et al*., 2025). However, the mechanisms underlying female sterility differ substantially. In the MDFS system, sterility results from somatic disruption of the female-specific *dsx* isoform, producing intersex females with impaired blood-feeding behaviour and consequently reduced malaria transmission (Kyrou *et al*., 2018; Strampelli *et al*., 2025). In contrast, disruption of *fle* occurs primarily within the germline, leaving external morphology, mating behaviour and blood-feeding largely unaffected. Consequently, sterile *fle^zpgp/+^* females are expected to remain capable of biting and potentially transmitting *Plasmodium*, an important distinction when considering operational deployment for malaria control.

This limitation may be overcome by combining *fle* targeting with promoters that exhibit broader expression, such as *vasa2*, as in the architecture of the MDFS system (Strampelli *et al*., 2025). Expanding Cas9 activity beyond the germline could couple disruption of oogenesis with female lethality, eliminating sterile but blood-feeding females and thereby reducing any residual contribution to parasite transmission. Whether such an approach can be achieved without compromising drive performance remains to be determined experimentally.

Collectively, our findings reveal a previously unrecognised role for *femaleless* in female gametogenesis and demonstrate that targeting genes essential for germline functions can prevent the spread and reduce the effectiveness of homing-suppression gene drives. Although this property limits the use of *fle* in self-sustaining population-suppression systems, it makes the gene an attractive target for male-driven, self-limiting strategies. More generally, our study highlights the importance of integrating developmental genetics with gene-drive engineering when identifying targets for next-generation vector-control technologies.

## Methods

### Generation of donor and CRISPR constructs

The *fle^DL^* docking line was created using a donor plasmid containing a GFP transcription unit (*3xP3::GFP*) flanked by two ΦC31 attP recombination sequences, inserted within the second RNA recognition motif (RRM) of the *fle* gene. We selected two gRNAs, 13051T14 (T14) and 13051T1 (T1), targeting highly conserved regions (T1 and T14 conservation scores are 0.434 and 0.344, respectively) within the second exon, spaced 74 bp apart (Fig. 1A). The gRNAs were chosen in highly conserved regions of the *Anopheles* species complex (Kranjc *et al*., 2021), including analysis of field populations of *A. gambiae* and *A. coluzzi* (Fig S1) (Clarkson *et al*., 2020; The Anopheles gambiae 1000 Genomes Consortium. Ag1000G phase 3 SNP data release. MalariaGEN, 2021).

We generated the donor plasmid Fle_DL_v2 (*pBac[AttB-3xP3::RFP-fle_homology-AttP-3xP3:GFP-AttP-fle_homology-AttB*]) using standard molecular cloning protocols. Briefly, we digested the MH541846 plasmid (Kyrou *et al*., 2018) with *Age*I and *Mlu*I to obtain the 5442 bp backbone fragment. The GFP marker cassette flanked by *ϕC31 attP* recombination sequences (1407 bp) was amplified from the same plasmid using primers GFP_gibson_filler (1398 R) – Rarm and GFP_gibson_filler (1 F) – Larm. Left and right homology arms (1023 bp and 1019 bp, respectively) were amplified from *A. gambiae* G3 wild-type genomic DNA using primers Larm_F/Larm_R and Rarm_F/Rarm_R (**Table S**), with KAPA HiFi HotStart ReadyMix PCR Kit (Kapa Biosystems). Fragments were assembled using Gibson assembly.

For the *zpg* multi-gRNA CRISPR construct (*pBac*[*AttB-U6::13051T14gRNA-3xP3:RFP-zpgp:hCas9-U6:13051T1gRNA-AttB*], we modified plasmid p17410 (Kyrou et al., 2018) which contained human-codon-optimised *Cas9* (*hCas9*) under control of the germline-specific *zero population growth* (*zpg*) promoter (*AGAP006241)* and a single U6:gRNA spacer-cloning cassette. The *13051T1 (T1)* gRNA was inserted by Golden Gate cloning after *Bsa*I digestion. To create the multiplex system, we amplified a second U6::gRNA cassette using primers containing *Pvu*I restriction sites, digested the *p17410_T1* gRNA plasmid with *Pvu*I, and inserted the second cassette. Finally, the *13051T14 (T14)* gRNA was inserted by Golden Gate cloning after *Bsa*I digestion.

For the *spo11* multi-gRNA CRISPR construct (*pBac[attB-U6::13051T14gRNA-3xP3::RFP-spo11p:hCas9-U6:13051T1gRNA-attB*]), we digested the *zpg* construct with *Age*I, *Sgs*I, and *Ade*I to isolate a 5968 bp fragment containing the *3xP3:RFP* marker, both *U6 gRNA* cassettes, and the two *attP* sites. The *spo11* promoter-*Cas9*-*spo11* terminator cassette was amplified from plasmid *gs24* (Grilli *et al*., 2026). The two fragments were assembled using Gibson assembly to produce the *pBF1* plasmid.

### Generation of the transgenic mosquito lines

For the *fle^DL^* docking line, we microinjected embryos from the *A. gambiae* G3 strain with the *Fle_DL_v2* donor plasmid (100 ng/μl) and a helper plasmid containing *zpg*-driven Cas9 and both gRNAs, T1 and T14 (100 ng/μl). G0 survivors were outcrossed to wild-type mosquitoes, and G1 progeny were screened for GFP-positive (GFP^+^) individuals using fluorescence microscopy.

The insertion site was validated by diagnostic PCR using primers WG_162 and GFPseqR, which bind within the GFP construct and the genomic flanking region, outside the homology arm sequence. To distinguish hemizygotes from homozygotes, we used primers WG_162 and WG_163 mapping outside the insertion site to amplify across the locus: hemizygotes yield both a 2045 bp amplicon (HDR event) and a 717 bp fragment (wild-type allele) while homozygotes produce only the 2045 bp band (Fig. S1). Sequencing of the amplicon confirmed the 72 bp deletion and its replacement with the GFP cassette.

For the *zpg*-*fle* gene drive lines (*fle^zpgp^*), we employed recombinase-mediated cassette exchange (RMCE). We microinjected *fle^DL^*embryos with the *zpg* CRISPR plasmid (250ng/μl) and a helper plasmid containing *vasa2*::*ΦC31* integrase (250ng/μl) (Bischof *et al*., 2007; Volohonsky *et al*., 2015). G0 survivors were outcrossed to wild-type individuals, and the G1 progeny were screened for RFP-positive (successful exchange) and GFP-negative (loss of original marker) individuals.

Successful cassette exchange and the orientation of the Cas9 coding sequence relative to *fle* were confirmed by diagnostic PCR using primers fleBF-1 through fleBF-7, binding within the construct and flanking genomic regions (Fig. S5). This approach yielded two independent lines with opposite Cas9 orientations: *fle^zpgp#5^*(Cas9 inverted relative to *fle*) and *fle^zpgp#9^* (Cas9 in the same orientation as *fle*). Both lines were maintained exclusively through males due to female sterility. After confirming similar phenotypes in both lines, we selected *fle^zpgp#5^*(renamed *fle^zpgp^*) for detailed mechanistic studies.

For *spo11*-fle gene drive lines, we followed the same RMCE strategy but injected the *spo11* CRISPR construct (BF1) into the *fle^DL^* embryos. Multiple independent lines were obtained, of which we retained *fle^spo11p#3^* and *fle^spo11p#6^* for characterisation based on robust establishment and differing Cas9 orientations. Most phenotypic analysis focused on *fle^spo11p#3^*

### Molecular validation and genotyping

Genomic DNA was extracted using the DNeasy Blood & Tissue Kit (Qiagen) PCR validation, or the Wizard Genomic DNA Purification kit (Promega) for amplicon sequencing. For assessment of genetic sex in *fle^DL/DL^* individuals, we used primers yob-F and yob-R to amplify a fragment to amplify a fragment of the Y chromosome-specific *yob* gene by PCR (Qiagen FastCycle) on genomic DNA from ten morphologically male *fle^DL/DL^* pupae. For assessment of genetic sex in *fle^DL/DL^* individuals, we used primers yob-F and yob-R

### Phenotypic assays: Survival and sex ratio analysis

To determine the developmental stage at which homozygous *fle^DL/DL^* females die, we conducted a survival assay using a Y-linked RFP marker to track genetic sex.

We crossed hemizygous *fle^DL/+^* females with males carrying both, the *Y^3xP3::RFP^* marker (Bernardini *et al*., 2014) and the *fle^DL/+^* allele (genotype *Y^3xP3::RFP^; fle^DL/+^*) (Supplementary Fig. S2). These males were generated by crossing *fle^DL/+^* females with *Y^3xP3::RFP^* males and selecting GFP-positive and RFP-positive male offspring. Fifty *Y^3xP3::RFP^; fle^DL/+^* males were backcrossed to fifty *fle^DL/+^* females to produce progeny that included wild-type, hemizygous and homozygous *fle^DL/+^* individuals of both genetic sexes (identifiable by RFP in males).

First instar larvae (L1) were screened using a Complex Object Parametric Analyser and Sorter (COPAS) to separate *fle^DL/+^* (intermediate GFP) from *fle^DL/DL^* (high GFP) individuals based on fluorescence intensity (Marois *et al*., 2012). Larvae were sorted into batches of 100 individuals, with at least three replicates per genotype (*fle^+/+^*, *fle^DL/+^* and *fle^DL/DL^*). Control batches consisted of L1 progeny from crosses between *Y^3xP3::RFP^* males and wild-type females. Larvae were reared under standard conditions and monitored daily. At pupal and adult stages, individuals were scored for sex (morphologically and by RFP expression) and GFP marker presence. Survival rates from L1 to adult were calculated for each genotype and sex combination.

### Fecundity and fertility assays

Individual fertility assays were conducted following established protocols (Kyrou *et al*., 2018; Fuchs *et al*., 2021), with modifications to account for the observed sterility phenotypes. For the *fle^DL^* transgene, at least 40 mosquitoes from each genotype of *fle^DL/+^* or *fle^DL/DL^* were crossed to an equal number of wild-type mosquitoes of the opposite sex and allowed to mate for 5 days. For gene drive lines (*fle^zpgp^* and *fle^spo11p^*), we crossed 50 hemizygous males or females with 50 wild-type individuals of the opposite sex. Wild-type x wild-type crosses (50 pairs) served as controls in all experiments.

After the mating period, females were blood-fed for 45 minutes. We did not pre-select blood-fed females; all females were transferred to individual oviposition cups two days post-blood meal, regardless of feeding status. Eggs were counted, and after hatching, larvae were counted and screened for fluorescent markers. For *fle^DL^* crosses involving homozygous males, all larvae were screened for GFP to determine parental genotype (homozygous produce 100% GFP-positive offspring). For gene drive crosses, RFP-positive larvae were counted as a proxy for drive inheritance, allowing calculation of homing rates. Females found dead in oviposition cups were included in the analysis if they had been alive at the time of transfer. Females that failed to lay eggs were dissected to check for sperm in the spermathecae. Statistical comparisons between genotypes were performed using unpaired t-tests or one-way ANOVA with Tukey’s post hoc test, as appropriate, when p < 0.05 considered significant.

The assess Cas9 contribution to sterility in *fle^zpgp#5/+^*females we employed AcrIIA4, an anti-CRISPR protein that specifically inhibits Cas9 activity. We crossed 30 *fle^zpgp#5/+^* males with 30 *vasa2::AcrIIA4 119* (GFP+) females (Taxiarchi *et al*., 2021). From the offspring, we selected the trans-hemizygous males and females and crossed them separately to wild-type counterparts. We also set crosses with the same number of individuals using *fle^zpgp#5/+^* males and females as comparisons. After 5-7 days of mating, the females were blood-fed for 45 minutes and after two days engorged 30 females from the *vasa2::AcrIIA4* crosses and 15 for *fle^zpgp#5/+^*crosses were transferred to individual oviposition cups. Eggs were counted, and after hatching, larvae were counted and screened for fluorescent markers. Homing rate for only 5 *fle^zpgp#5/+^*males was scored.

### Bipartite system crosses

To confirm that female sterility in the *fle^zpgp^* resulted from loss of fle function rather than transgene position effects, we tested a bipartite system with separated Cas9 and gRNA components. We crossed males from the *zpgp:Cas9-2L-20B* (*zpgp:Cas9*) line (carrying zpg-driven Cas9 at an out-of-locus insertion site) with females from the *fle^gRNAs^*line (carrying the T1 and T14 gRNAs in a piggyBac insertion). From the F1 progeny, we selected more than 30 trans-hemizygous males (carrying both transgenes) and more than 30 trans-hemizygous females and crossed each sex to wild-type mosquitoes of the opposite sex. Individual fertility assays were conducted as described above, with at least 26 trans-hemizygous females tested.

### Crosses to assess Cas9 activity and paternal deposition

To distinguish between paternal CA9 deposition and somatic Cas9 expression, we designed crosses using the *fle^DL/+^* as a resistant target. We crossed fifty *fle^zpgpp/+^* males with fifty *fle^DL/+^* females to generate *fle^zpgp/DL^* sons (RFP-positive and GFP-positive). These males inherit the gene drive (RFP-positive) and the docking allele (GFP-positive), with *fle^DL^* serving as a resistant allele due to its large marker cassette inserted, which prevents gRNA binding. Fifty *fle^zpgp/DL^* males were then crossed to fifty wild-type females, alongside a control cage of fifty wild-type females crossed to fifty wild-type males.

After five days of mating, females were blood-fed and allowed to oviposit. Hatched larvae were divided into trays of 200 individuals. At the pupal stage, we screened for fluorescent markers and selected 2 x 60 individuals of each genotype, *fle^zpgp^*^/+^ (RFP-positive and GFP-negative) and *fle^DL^/+* (RFP-negative and GFP-positive) and each sex for genomic DNA extraction and amplicon sequencing. In the offspring that did not inherit *fle^zpgp^*^/+^ (RFP-negative), modifications of the maternally-derived wild-type allele could only result from paternally deposited Cas9, while modifications in RFP-positive offspring indicate Cas9 activity in those individuals.

### Tissue-specific Cas9 activity analysis

To determine whether Cas9 activity occurred in somatic tissues in addition to the germline, we dissected 20 heterozygous *fle^zpgp^/+* females and 20 wild-type females that had not been blood-fed. Ovaries (excluding spermathecae) and remaining body tissue (carcasses), were stored separately at −20°C in 180 µL of ATL buffer (DNeasy Blood and Tissue Kit, QIAGEN) prior to genomic DNA extraction. Amplicon sequencing across the *fle* target region was performed separately for ovarian and somatic tissues, with mutation frequencies compared between tissue types and to wild-type controls.

### Crosses and analysis for *spo11* promoter system

For initial characterisation of *spo11*-promoter timing effects, we crossed 50 females of genotype *spo11p:Cas9a* (an out-of-locus GFP-positive transgene mapping to chromosome 2L20D) with 41 males of genotype *gRNAs^fle^* (RFP-positive, piggyBac insertion maintained in homozygosity, containing both T1 and T14 gRNAs under U6 promoters). From the F1 progeny, we selected 50 trans-hemizygous females (*spo11p:Cas9a*/+; *gRNAs^fle^*/+, expressing both RFP and GFP) and crossed them to 50 wild type males. The reciprocal cross (trans-hemizygous males crossed to wild-type females) was also performed. F2 progeny from both crosses were collected for amplicon sequencing of the *fle* target site.

For the integrated *fle^spo11p^* gene drive lines, individual fertility assays were conducted as described above, with 30 hemizygous *fle^spo11p/+^* individuals of each sex crossed to wild-type. Progeny were screened for RFP to calculate transmission rates, and separate batches were collected for amplicon sequencing from both male and female parents.

To assess homing events in the germline of *fle^spo11p/+^*females we used the *dsxF^-^* (*dsxDL 4050E5*; GFP*+*) line (Kyrou *et al*., 2018) bearing a GFP marker on the homologous chromosome inserted in the female exon of *dsx*. We crossed 50 *fle^spo11p/+^*males to 50 *dsxF^-^* females. From the offspring, we selected 30 trans-hemizygous *fle^spo11p^/ dsxF^-^* males and females and crossed them separately to wild-type counterparts. After 5-7 days of mating, the females were blood-fed for 45 minutes and after two days an egg bowl was introduced in the cage to collect embryos. Embryos were transferred to three trays for the male *fle^spo11p^/ dsxF^-^* progeny and to a single tray for the female reduced progeny. Larvae were then counted and manually sorted for fluorescent markers.

### Ovarian dissection and developmental stage analysis

To determine when Cas9-mediated mutagenesis occurs in *fle^spo11p/+^*females and whether mutated oocytes are eliminated, we dissected ovaries at a specific developmental timepoint when *spo11* expression peaks. Mated *fle^spo11p/+^*females (n = 20) and wild type controls (n = 20) were blood-fed and dissected 3 hours post-blood meal, a timepoint when spo11-driven Cas9 expression is increasing (Terradas *et al*., 2021). Ovaries were carefully separated from spermathecae (which were discarded) and immediately stored at −20 °C in ATL buffer (DNeasy Blood and Tissue Kit, QIAGEN) for genomic extraction and amplicon sequencing.

For females that failed to oviposit, we performed dissections at 5 days post-blood meal to assess ovarian development. Ovaries were examined under a dissecting microscope, and representative samples were photographed.

### Amplicon sequencing and mutation analysis

Genomic DNA was extracted using the Wizard Genomic DNA Purification kit (Promega) for large samples or the DNeasy Blood and Tissue Kit (QIAGEN) for small tissue samples. All samples were homogenised using the Precellys Evolution Homogeniser prior to DNA extraction.

To analyse drive transmission and resistance alleles in progeny, we used the following approaches: i. For the *fle^zpgp^/fle^DL^* x wild-type crosses: We collected two samples of 60 individuals each, for both genotypes (*fle^zpgp/+^*, RFP-positive and *fle^DL/+^*, GFP-positive), and extracted DNA; ii. For RFP-negative progeny (resistance allele analysis): We crossed 100 *fle^zpgp^* heterozygous males with 150 wild-type females and used a COPAS FlowPilot Cytometer (UNION BIOMETRICA) to sort L1 larvae by RFP fluorescence. We isolated 59 RFP-negative larvae from more than 4500 screened L1 larvae and pooled them for DNA extraction.

For *fle^spo11p/+^* RFP-negative progeny (resistance allele analysis), we crossed 50 *fle^spo11p/+^* heterozygous males to 50 wild-type females and vice versa and manually sorted pupae by RFP fluorescence. Similar crosses were performed for *fle^spo11p#3^* and *fle^spo11p#6^* lines. RFP-negative pupae were collected in pools, homogenised in NLS buffer (Promega), DNA extracted, and sent for amplicon sequencing. DNA from RFP-negative progeny of *fle^spo11p#3/+^*fathers was extracted separately from 3 different collections of pupae (total = 40 individuals), pooled, and then sent for amplicon sequencing. To assess Cas9 activity in *fle^spo11p/+^*females, we dissected the ovaries from 20 individuals and used wild-type females as controls. Ovaries were dissected in a drop of cold 1X PBS and collected in ATL buffer before DNA extraction.

For amplicon sequencing of F1 embryos, 100 *fle^spo11p#3/+^*females were crossed with an equal number of wild-type males. A control was set up with a wild-type cross of the same size. Females were mated for three days and blood-fed twice, at 3 and 6 days PBM, to collect two batches of embryos. 24-hour-old embryos were collected overnight in an egg bowl containing filter paper and saline, then transferred to homogenisation tubes using Flystuff (57-103) Nitex Nylon Mesh and Flystuff (46-102) Mesh Basket. DNA was extracted using the Wizard Genomic DNA Purification kit (Promega).

For all samples, we amplified a 476 bp region spanning both *fle* target sites using primers *fle_amplicon2F* and *fle_amplicon2R*, with KAPA HiFi HotStart ReadyMix kit for 25 cycles. Amplicons were purified using the QIAquick PCR purification kit (QIAGEN) or Wizard SV Gel and PCR Clean-Up system (Promega) and submitted to Genewiz for Illumina sequencing. Sequencing data were analysed using CRISPResso2 (Clement *et al*., 2019) to quantify modification frequencies, indel sizes, and mutation classes. For comparisons between samples, we calculated the percentage of reads carrying modifications relative to the total number of high-quality reads, with wild-type controls run in parallel to establish background mutation rates.

### Phenotypic characterisation and confocal microscopy

For external morphology and spermatheca examination, mosquitoes were anaesthetised on ice for 5 minutes and dissected in 1X PBS (pH 7.4). Ovaries were transferred to a drop of PBS on a Petri dish and imaged using an Olympus MVX10 microscope. Spermathecae were visualised directly using an EVOS XL Core Cell imaging system to assess sperm presence and abundance. For quantification, we scored spermathecae as containing sperm (regardless of quantity) or empty and noted any abnormalities in spermatheca number or morphology.

For Confocal imaging of ovarian structures, ovaries were dissected from anaesthetised mosquitoes, collected in cold PBS on ice, and fixed in 4.5% formaldehyde in PBS with 0.1% Triton-X for 30 minutes at room temperature on a nutator. Samples were washed three times for 10 minutes in PBS- T (PBS with 0.1% Triton-X), then mounted in VECTASHIELD® PLUS antifade mounting medium with DAPI (H-2000). Images were captured using either a Leica CF6-SP8 inverted confocal microscope or a Leica LS1-Stellaris 5 laser scanning confocal microscope, both equipped with 63X oil-immersion objectives. Z-stacks were acquired at optimal intervals determined by the Leica imaging software.

Image processing was performed using Leica Application Suite X (LAS X), Imaris Viewer 10.2.0, Fiji (ImageJ2 2.14.0/1.54f) and Adobe Photoshop 26.8.1. For maximum-intensity projections, confocal z-stacks were processed in Fiji or Imaris Viewer. Brightness and contrast adjustments were applied uniformly across entire images using Adobe Photoshop and Lightroom 2025, and images were cropped as needed for figure presentation. No manipulations were performed that would alter the interpretation of the biological structures visualised.

### Mathematical modelling

To estimate the level of suppression in a wild population following the release of a homing construct we use a population dynamics and genetics model following that of Burt and Deredec. We model a single, well-mixed population of infinite size using a deterministic model in discrete time with non-overlapping generations. We allow for two sexes (male and female) and assume density-dependent mortality occurs during the juvenile stage according to the Beverton-Holt model and assume the population had an R_m_ of 6. We model a single locus with four possible alleles: wild-type (A1), genetic construct (A2) and two products of non-homologous end joining repair. The first product (A3) is resistant to cleavage at only one of the two target sites, allowing cleavage to still occur in A2/A3 heterozygotes. The second product (A4) is resistant to cleavage at both target sites, preventing homing entirely. All disrupted loci (A2, A3 and A4) cause recessive larval lethality. Females homozygous for any combination of disrupted alleles (lacking A1 entirely) die as larvae before density-dependent mortality occurs, whilst heterozygous females carrying at least one A1 allele mature as fully fertile adults. Based on observed phenotypes of the fle^GD#5^, the genetic construct is assumed to cause sterility specifically in A1/A2 heterozygous females. All males are assumed to have equal fitness to the WT males. Sterility is modelled by assuming that adult females can bite but do not produce offspring. Table 1 summarises the fitness costs.

**Table 1.** Genotype-specific fitness in females.

| Genotype | Fitness |  |
| --- | --- | --- |
|  | Larval survival | Fertility |
| A1/A1 | 1 | 1 |
| A1/A2 | 1 | 0 |
| A1/A3 | 1 | 1 |
| A1/A4 | 1 | 1 |
| A2/A2 | 0 | 1 |
| A2/A3 | 0 | 0 |
| A2/A4 | 0 | 1 |
| A3/A3 | 0 | 1 |
| A3/A4 | 0 | 1 |
| A4/A4 | 0 | 1 |

To model the impact of the construct allele on gamete production, we assumed that A1/A2 heterozygotes, the WT allele was cleaved with probability *c_X_*, after which the cleaved allele could be repaired via NHEJ with probability *j_x_* or via homology-directed repair to form an A21 − *j_X_*with probability ￼*X*, where *X* = *M* denotes the rates in males (*X* = *F*) or females (￼). NHEJ products are either A4 or A3, with probability *p_x_* or 1 − *p_x_* respectively. The probability of each gamete being produced from each parent is shown in Table 2 and a summary of the parameters used is in Table 3.

**Table 2.** Gamete table.

| Parental genotype | A1 | A2 | A3 | A4 |
| --- | --- | --- | --- | --- |
| A1A1 | 1 | 0 | 0 | 0 |
| A1A2 | $\frac{1 - c_x}{2}$ | $\frac{1 + c_x(1 - j_x)}{2}$ | $\frac{c_x j_x(1 - p_x)}{2}$ | $\frac{c_x j_x(1 - p_x)}{2}$ |
| A1A3 | $\frac{1}{2}$ | 0 | $\frac{1}{2}$ | 0 |
| A1A4 | $\frac{1}{2}$ | 0 | 0 | $\frac{1}{2}$ |
| A2A2 | 0 | 1 | 0 | 0 |
| A2A3 | 0 | $\frac{1 + c_x(1 - j_x)}{2}$ | $\frac{1 - c_x}{2}$ | $\frac{c_x j_x}{2}$ |
| A2A4 | 0 | $\frac{1}{2}$ | 0 | $\frac{1}{2}$ |
| A3A3 | 0 | 0 | 1 | 0 |
| **A3A4** | 0 | 0 | $\frac{1}{2}$ | $\frac{1}{2}$ |
| **A4A4** | 0 | 0 | 0 | 1 |

**Table 3.** Inheritance parameters.

| Parameter | Description | Data Source | Parameter |
| --- | --- | --- | --- |
| **d** | Proportion of offspring carrying the mutation | Experiment | 0.980 |
| **e** | Probability of homing | $2d - 1$ | 0.960 |
| **j** | Probability of NHEJ given cleavage (c) | $\frac{(1 - e)u}{e + (1 - e)u}$ | 0.036 |
| **u** | Proportion of non-homed chromosomes which are NHEJ | Experiment | 0.894 |
| **c** | Probability of cleavage | $e + (1 - e)u$ | 0.997 |
| **p** | Probability of a WT allele being converted to a fully cleavage resistant allele use to NHEJ. | Experiment | 0.803 |

### Statistical Analysis

Data are presented as mean ± standard error of the mean (s.e.m.) unless otherwise stated. Sample sizes are indicated in the text or figure legends. For fertility assays, we compared larval output and hatching rates between genotypes using unpaired t-tests. (for two-group comparisons) or one-way ANOVA with Tukey’s post-hoc test (for multiple group comparisons). For proportions (e.g., transmission rates, mutation frequencies), we report percentages with s.e.m. calculated from replicate crosses or samples. Statistical significance was set at p < 0.05, with exact p-values reported when p < 0.05. Analysis was performed using GraphPad Prism v10. No data were excluded from analysis.

## Supplementary figures

**Supplementary Fig. S1.**
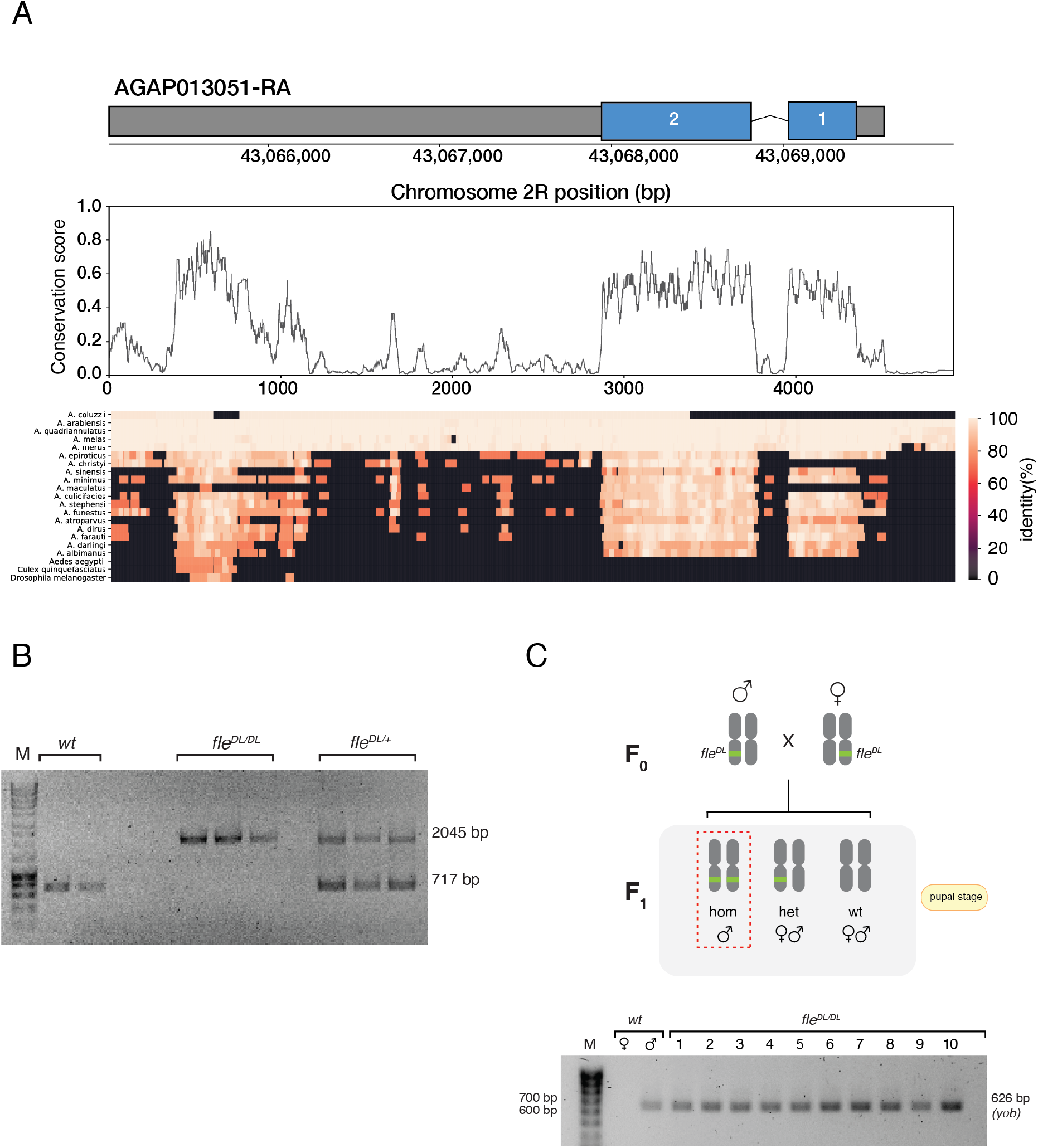
Target site conservation and validation of fle^DL^ integration and genotype. (**A**) Conservation analysis of the *femaleless* (*fle*, AGAP013051) locus across 21 *Anopheles* species. Gene structure indicating exon positions. Middle, nucleotide conservation scores across the locus. Bottom, percentage identity heatmap across species. gRNA target sites T1 and T14 are located within highly conserved regions (>90% identity) selected to minimise the formation of resistant alleles. (**B**) Genotyping PCR confirming correct integration of the GFP marker cassette. Homozygous *fle^DL/DL^* individuals show a single 2045 bp amplicon corresponding to the integrated allele (primers WG162/WG163); hemizygous *fle^DL/+^* individuals show both the 2045 bp integration band and a 717 bp wild-type band representing the wild-type (*wt)* allele without integration; wild-type (*wt*) individuals show only the 717 bp band. M, 1 kb DNA ladder. (**C**) Genetic cross and sex genotype validation. Top, schematic of crosses between *fle^DL/+^*males and females (F_0_) and resulting F_1_ progeny genotypes. All homozygous pupae are phenotypically male (red dashed outline). Bottom, PCR amplification of the Y-linked *yob* gene confirms that all morphologically male *fle^DL/DL^* individuals (n = 10) carry the Y chromosome, indicating that homozygous females are absent owing to early larval lethality. Amplicon sizes and marker bands are indicated.

**Supplementary Fig. S2.**
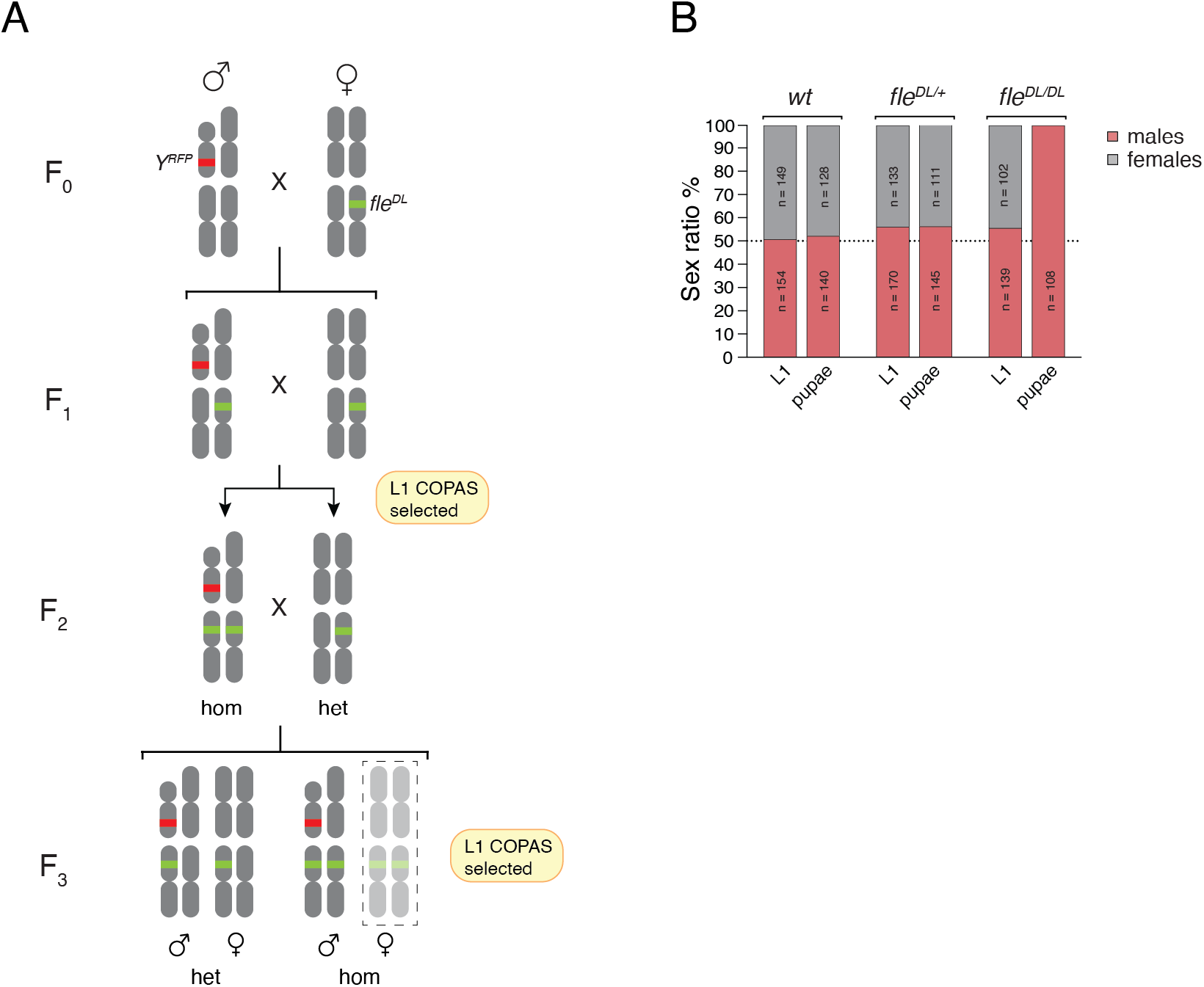
Homozygous *fle^DL^* females exhibit larval-stage lethality. (**A**) Crossing scheme using Y-linked RFP marker to track genetic sex. F_0_ males carrying *Y^RFP^* were crossed to *fle^DL/+^*females. In the F_1_ generation, *fle^DL/+^* males carrying *Y^RFP^* were selected and crossed to *fle^DL/+^*females. F_2_ progeny comprising homozygous (*fle^DL/DL^*) and heterozygous (*fle^DL/+^*) individuals were identified by GFP intensity at the L1 stage using COPAS sorting and intercrossed. To enrich for homozygotes, F_2_ *fle^DL/DL^* males were crossed to *fle^DL/+^* females, and F_3_ progeny were similarly sorted and analysed. (**B**). Sex ratio at the first larval (L1) and pupal stages for each genotype. Wild-type and *fle^DL/+^* individuals display an approximately equal sex ratio at both stages. In contrast, *fle^DL/DL^* individuals show a balanced sex ratio at L1 but are exclusively phenotypically male at the pupal stage, with genetically female (RFP-negative) individuals absent, indicating female-specific lethality during larval development (L2-L3 stages).

**Supplementary Fig. S3.**
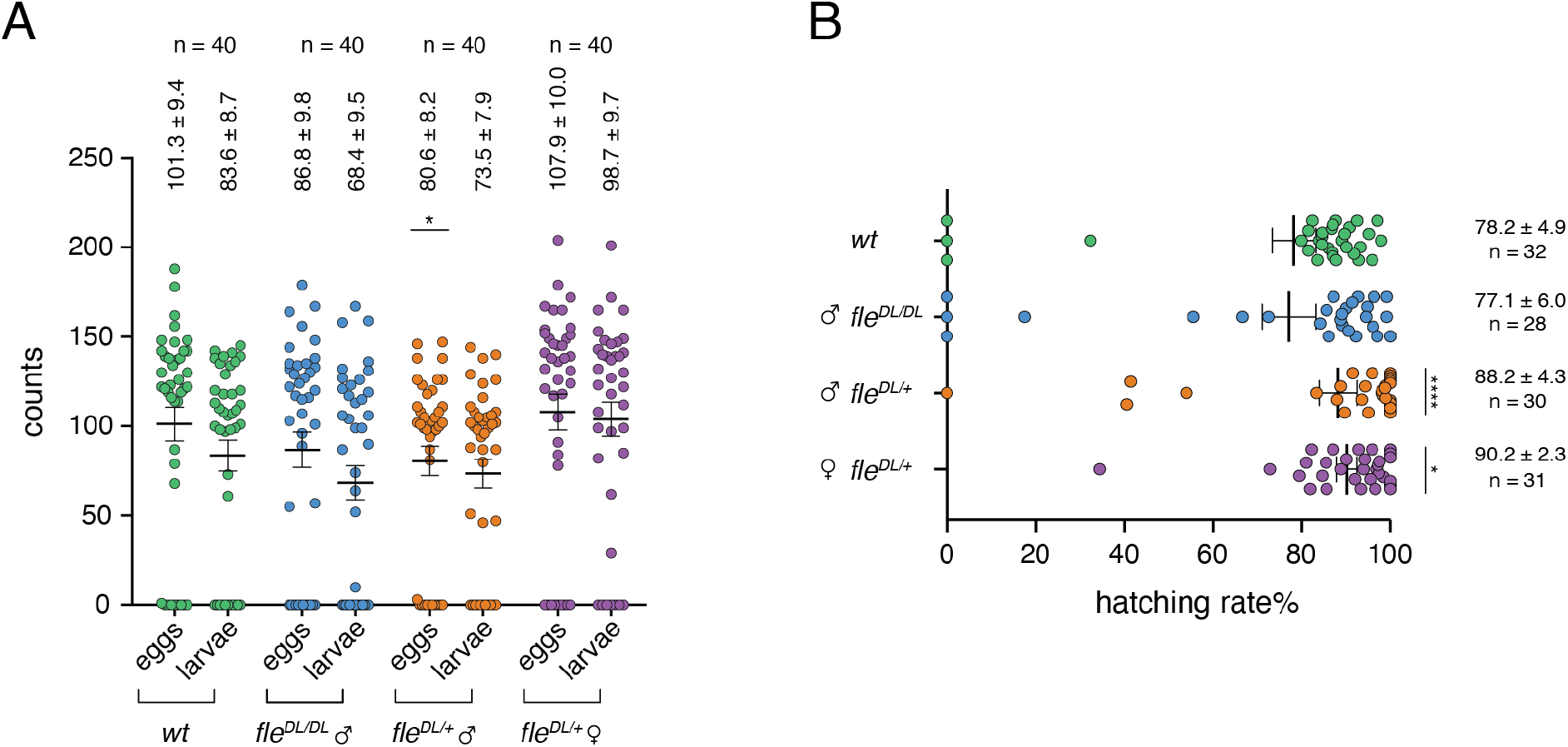
Fertility assays of *fle^DL^* hemizygous and homozygous mosquitoes crossed to wild-type. (**A**) Egg production and larval hatching across *fle^DL^*crosses. Egg output and larval hatching from individual layings across three cross types (left to right): wild-type (*wt*), wild-type females mated to *fle^DL/DL^*, wild-type females mated to *fle^DL/+^*and *fle^DL/+^* females crossed to wild-type. Note that *fle^DL/DL^*females were excluded from analysis owing to lethality at larval stages. Each dot represents an individual laying (n = 40 per group). Horizontal lines indicate mean ± s.e.m. with values shown above each group. Eggs output was compared with wild-type using a Kruskal-Wallis test with Dunn multiple-comparisons correction, revealing a significant difference for crosses involving *fle^DL/+^* male (P = 0.0387). No significant differences were observed in larval output relative to wild-type (Dunn multiple-comparisons *fle^DL/DL^* males, P = 0.8443; *fle^DL/+^* males, P = 0.6495; *fle^DL/+^* females, P = 0.3734). (**B**) Larval hatching efficiency in *fle^DL^* crosses. Percentage larval output (hatching rate) from crosses between wild-type (wt) homozygous *fle^DL/DL^* males, and hemizygous *fle^DL^* males and females crossed to wild-type. Each dot represents an individual deposition with at least one egg; horizontal lines indicate mean ± s.e.m., with values shown to the right of each group. Statistical analysis was performed using a Kruskal-Wallis test with Dunn’s multiple comparisons correction. Compared with wild-type, larval hatching was significantly increased in crosses involving *fle^DL/+^* males (P < 0.0001) and *fle^DL/+^* females (P = 0.01620), but not in crosses involving *fle^DL/DL^* males (P = 0.6271). Asterisks indicate statistical significance.

**Supplementary Fig. S4.**
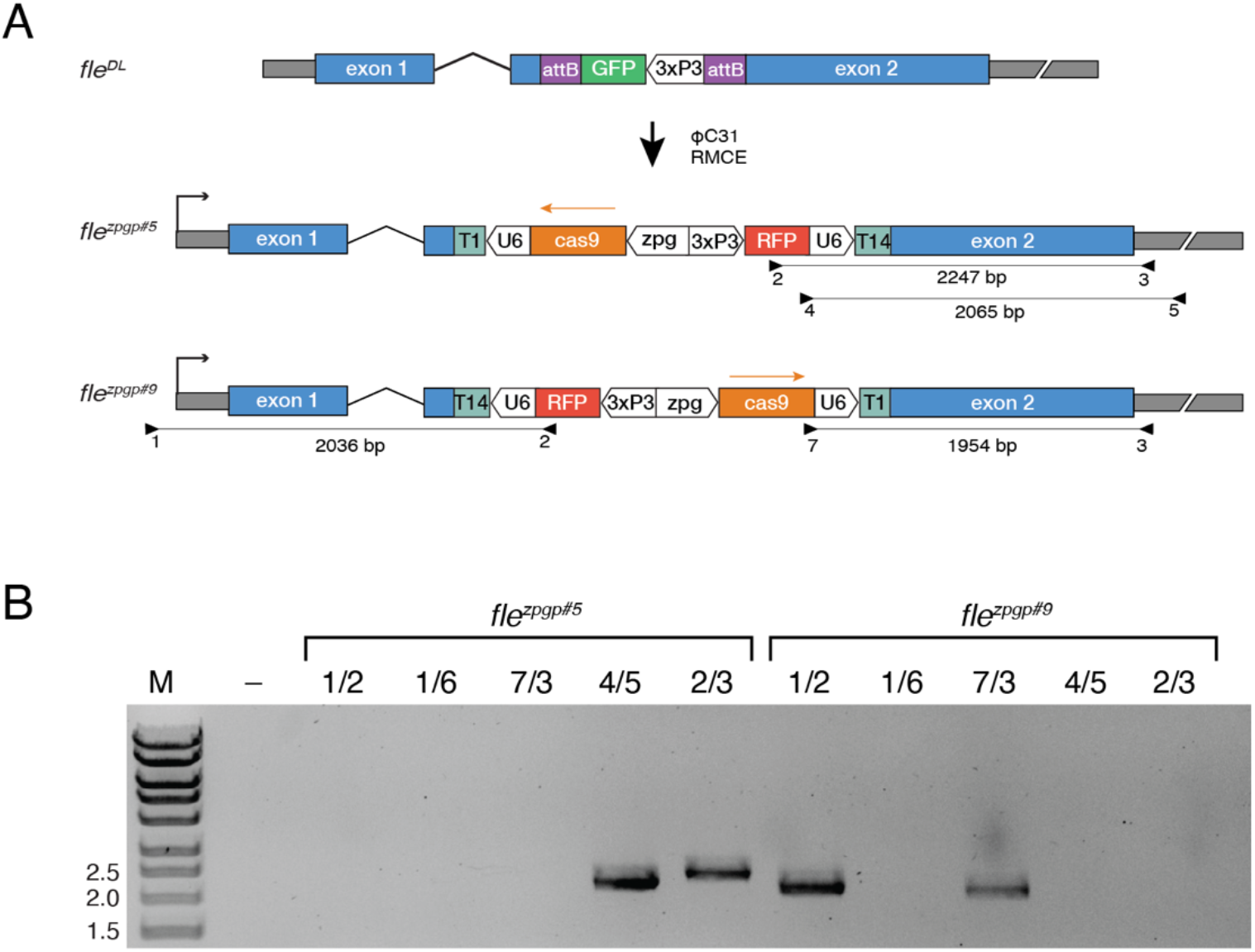
Orientation of the *fle^zpgp^* cassette within the *fle* gene. **(A)** Schematic representation of recombinase-mediated cassette exchange at the *fle* locus showing two possible orientations of the integrated *fle^zpgp^* cassette. In one configuration (*fle^zpgp#5^*), *cas9* is oriented opposite to the *fle* gene; in the alternative configuration (*fle^zpgp#9^*), *cas9* is aligned in the same orientation as *fle*. Primer positions are indicated by numbered arrows and expected amplicon sizes for diagnostic PCR are shown beneath each construct. **(B)** Diagnostic PCR of genomic DNA from *fle^zpgp#5^* (lanes 2-6) and *fle^zpgp#9^* (lanes 7-11) using the indicated primer pairs. The observed banding patterns confirm the cassette integration orientation in each transgenic line. M, DNA size marker. fleBF-1 to -7 primers in Supplementary Table S8.

**Supplementary Fig. S5.**
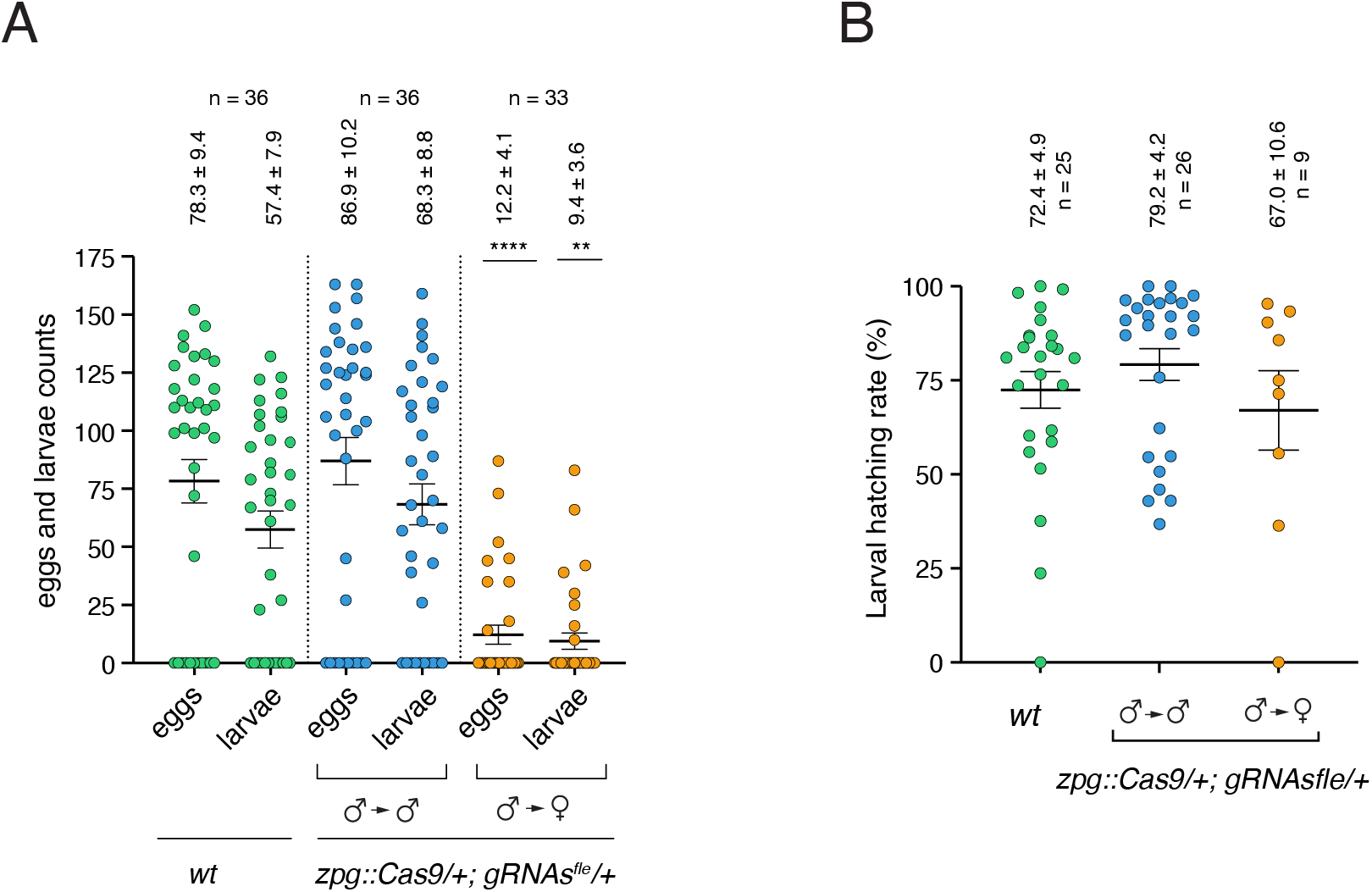
Fecundity and fertility of mosquitoes carrying *fle* mutations induced by *zpg::cas9* and *gRNA*s. **(A)** Egg production and larval output from individual layings across three cross types (left to right): wild-type controls (*wt*), wild-type females crossed to *zpg::Cas9; gRNAs^fle^*males and *zpg::Cas9; gRNAs^fle^* females crossed to wild-type males. Progeny derived from *zpg::Cas9* fathers are indicated ♂→♂ for males and ♂→♀ for females. Each dot represents an individual laying; horizontal bars indicate mean ± s.e.m. with values shown above each group, and n denotes the number of depositions. Compared with wild-type, *zpg::Cas9; gRNAs^fle^* females show a significant reduction in both eggs and larval output (P < 0.0001 and P = 0.0025, respectively), whereas *zpg::Cas9; gRNAs^fle^* males do not differ from control (P > 0.9999; Kruskal-Wallis test with Dunn’s multiple comparisons correction). (**B**) Larval hatching rates for the same crosses. Each dot represents an individual deposition; horizontal bars indicate mean ± s.e.m. with values shown above each group. Despite reduced fecundity, *zpg::Cas9; gRNAs^fle^* females show no significant difference in hatching rates compared to wild-type or male crosses (P = 0.2076, Kruskal-Wallis test).

**Supplementary Fig. S6.**
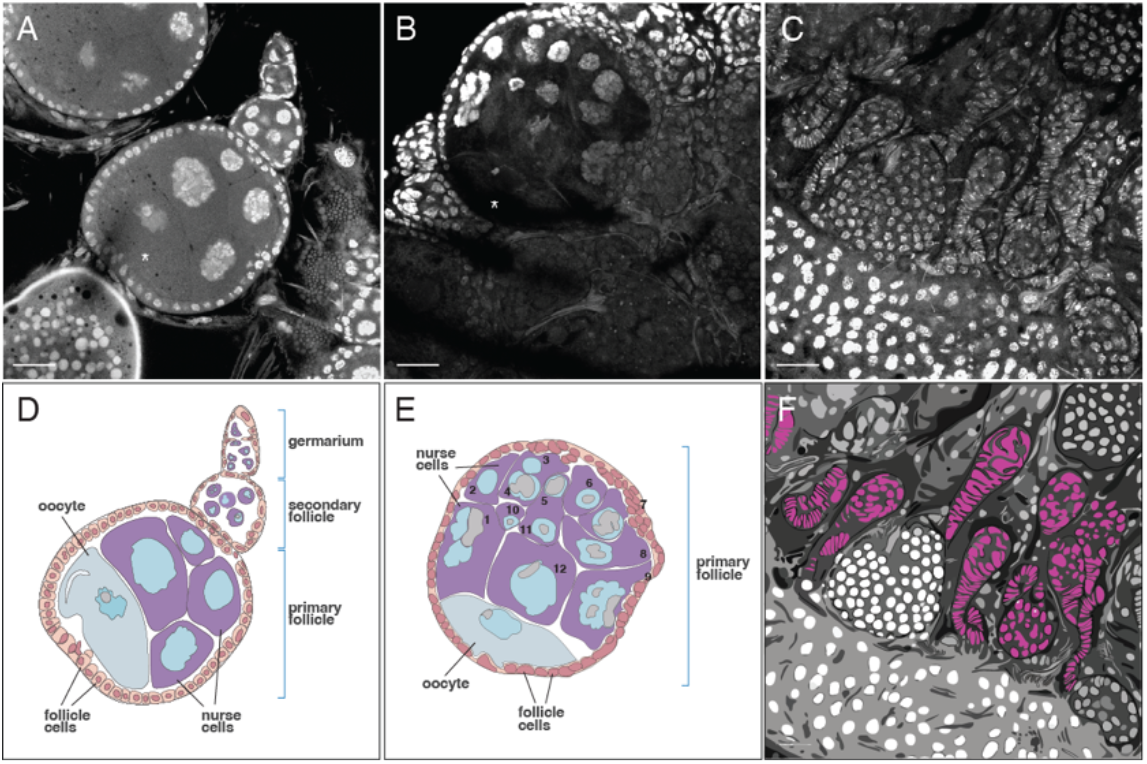
**Ovarian defects underlying sterility in *fle^zpgp^* females.**(**A**-**C**) Representative high-magnification confocal images of DAPI-stained ovaries from *fle^DL/+^* (**A**) and *fle^zpgp#5/+^*females (**B**-**C**). (**A**) Normal ovariole structure in *fle^DL/+^*, showing germarium, secondary and primary follicles with the typical complement of nurse cells. (**B**) *fle^zpgp#5/+^* follicle displaying an abnormal number of nurse cells compared with wild-type. (**C**) *fle^zpgp#5/+^* ovarioles exhibiting increased stacks of stalk cells. Oocytes are indicated by asterisks. (**D**-**F**) Schematic representation of corresponding follicle structures in *fle^DL/+^* (**D**) and *fle^zpgp#5/+^* ovaries (**E**-**F**). (**E**) *fle^zpgp#5/+^* follicles contain excess nurse cells (numbered 1-12), exceeding the normal complement of 7. (**F**) Abnormal accumulation of stalk cell structures in *fle^zpgp#5/+^* ovarioles. Scale bar: 20 μm.

**Supplementary Fig. S8.**
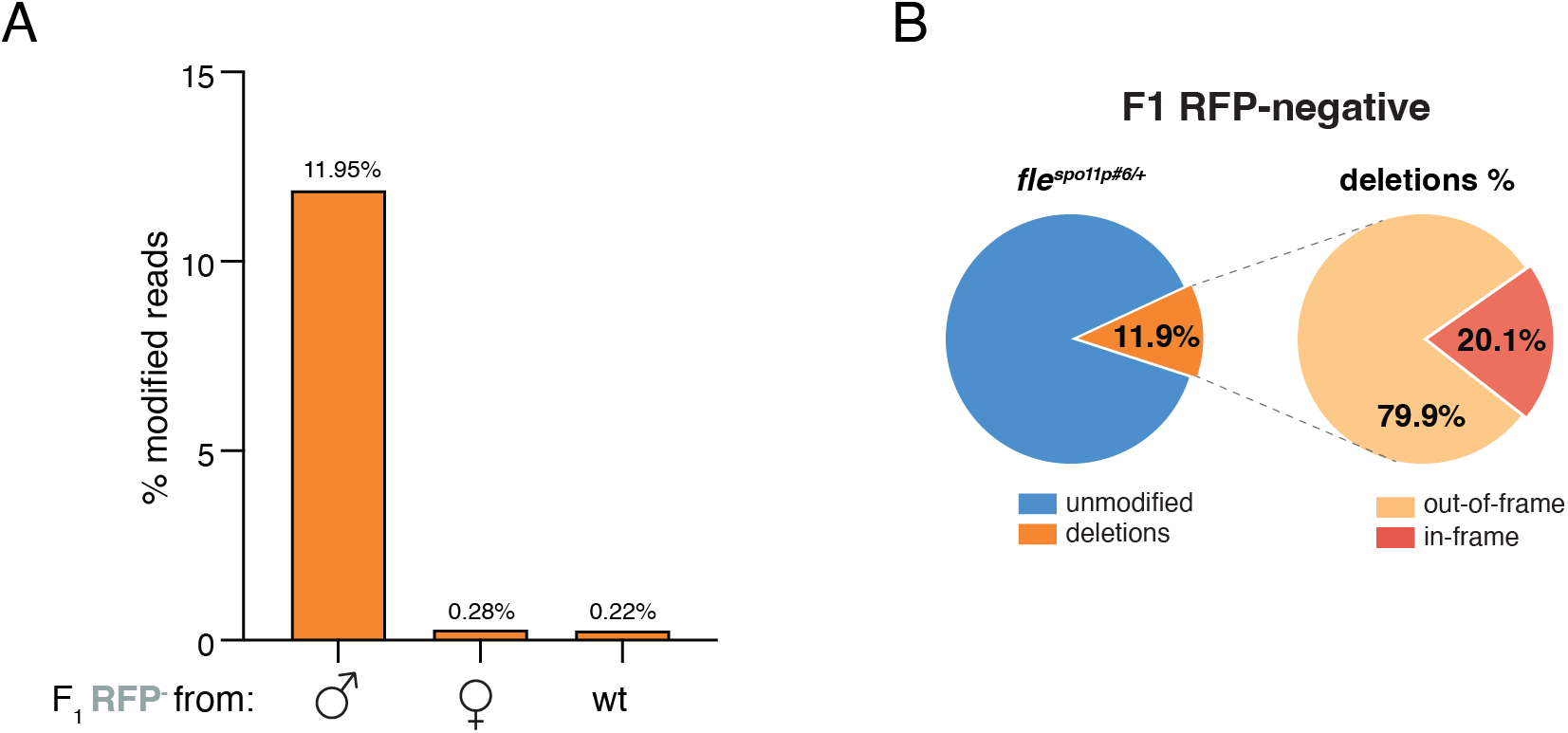
Mutation spectrum and frequency in *fle^spo11p#6/+^* non-drive progeny. (**A**) Percentage of modified reads from amplicon sequencing of F1 RFP-negative progeny derived from *fle^spo11p#6/+^* fathers, *fle^spo11p#6/+^*mothers and wild-type controls. Elevated modification rates are observed in progeny from *fle^spo11p#6/+^* fathers (11.95%) compared to maternal inheritance (0.28%) and wild-type control (0.22%) (**B**) Left: proportion of modified and unmodified reads in RFP-negative progeny from *fle^spo11p#6/+^* fathers, showing that 11.9% of reads carry mutations. Right: breakdown of mutation types among modified reads with a predominance of frameshift deletions (79.9%) over in-frame deletions (20.1%).

**Supplementary Fig. S9.**
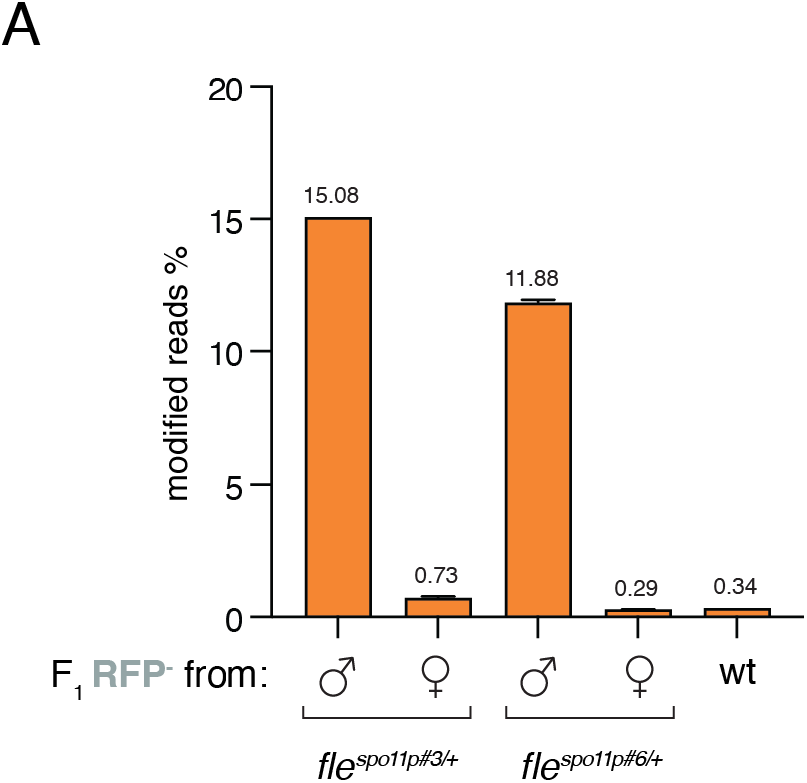
Reproducible sex biased mutation inheritance across independent *fle^spo11p^* lines. (**A**) Percentages of modified reads detected by amplicon sequencing in F_1_ RFP-negative progeny from *fle^spo11p#3^*and *fle^spo11p#6^* individuals crossed to wild-type. Progeny from males show elevated mutation frequencies (*fle^spo11p#3^*, 15.08%, n = 40; *fle^spo11p#6^*, 11.88%, n = 96), whereas progeny from females display low mutation rates (*fle^spo11p#3^*, 0.73%, n = 110) and (*fle^spo11p#6^*, 0.28%, n = 87) comparable to the wild-type control (0.34%). Error bars represent s.e.m. The consistent male-female asymmetry observed across independent lines indicates reproducible transmission rates.

## Acknowledgments

We thank Alekos Simoni and John Connolly for their valuable feedback. This work was supported in whole or in part by the Bill & Melinda Gates Foundation [Grant Number INV006610 “Target Malaria Phase II”] and Open Philanthropy (OPP1210755) (A.B. and A.C.).

## Author contributions

B.F. and W.G. conceived the project; B.F., W.G. and F.B. designed the research; B.F., W.G., J.P. and L.M. performed research; B.F. performed visualisation; B.F. supervised the project; N.K. and K.W. performed computational analysis; A.S. supplied reagents; B.F. wrote the original draft; B.F. and F.B. wrote, reviewed and edited the final manuscript with input from all authors; A.C. and A.B. acquired funding.

## Additional information

Supplementary data to be added

**Correspondence** and requests for materials should be addressed to Barbara Fasulo or Federica Bernardini

